# High-throughput inference of pairwise coalescence times identifies signals of selection and enriched disease heritability

**DOI:** 10.1101/276931

**Authors:** Pier Francesco Palamara, Jonathan Terhorst, Yun S. Song, Alkes L. Price

**Affiliations:** Department of Statistics, University of Oxford, Oxford, UK; Department of Epidemiology, Department of Biostatistics, Harvard T.H. Chan School of Public Health, Boston, MA, USA; Program in Medical and Population Genetics, Broad Institute of MIT and Harvard, Cambridge, MA, USA; Department of Statistics, University of Michigan, Ann Arbor, MI, USA; Department of Statistics, Computer Science Division, University of California, Berkeley, Berkeley, CA, USA; Chan Zuckerberg Biohub, San Francisco, CA 94158, USA

## Abstract

Interest in reconstructing demographic histories has motivated the development of methods to estimate locus-specific pairwise coalescence times from whole-genome sequence data. We developed a new method, ASMC, that can estimate coalescence times using only SNP array data, and is 2-4 orders of magnitude faster than previous methods when sequencing data are available. We were thus able to apply ASMC to 113,851 phased British samples from the UK Biobank, aiming to detect recent positive selection by identifying loci with unusually high density of very recent coalescence times. We detected 12 genome-wide significant signals, including 6 loci with previous evidence of positive selection and 6 novel loci, consistent with coalescent simulations showing that our approach is well-powered to detect recent positive selection. We also applied ASMC to sequencing data from 498 Dutch individuals (Genome of the Netherlands data set) to detect background selection at deeper time scales. We observed highly significant correlations between average coalescence time inferred by ASMC and other measures of background selection. We investigated whether this signal translated into an enrichment in disease and complex trait heritability by analyzing summary association statistics from 20 independent diseases and complex traits (average *N*=86k) using stratified LD score regression. Our background selection annotation based on average coalescence time was strongly enriched for heritability (p = 7×10^−153^) in a joint analysis conditioned on a broad set of functional annotations (including other background selection annotations), meta-analyzed across traits; SNPs in the top 20% of our annotation were 3.8x enriched for heritability compared to the bottom 20%. These results underscore the widespread effects of background selection on disease and complex trait heritability.

## Introduction

Recently developed methods such as the Pairwise Sequentially Markovian Coalescent (PSMC)^1^ utilize Hidden Markov Models (HMM) to estimate the coalescence time of two homologous chromosomes at each position in the genome^1-6^, leveraging previous advances in coalescent theory^7-11^. These methods have been broadly applied to reconstructing demographic histories of human populations^12-20^. More generally, methods for inferring ancestral relationships among individuals have potential applications to detecting signatures of natural selection^21^, genome-wide association studies^22-24^, and genotype calling and imputation^25-28^. However, all currently available methods for inferring pairwise coalescence times require whole genome sequencing (WGS) data, and can only be applied to small data sets due to their computational requirements.

Here, we introduce a new method, the Ascertained Sequentially Markovian Coalescent (ASMC), that can efficiently estimate locus-specific coalescence times for pairs of chromosomes using only ascertained SNP array data, which are widely available for hundreds of thousands of samples^29^. We verified ASMC’s accuracy using coalescent simulations, and determined that it is orders of magnitude faster than other methods when WGS data are available. Leveraging ASMC’s speed, we analyzed SNP array and WGS data sets with the goal of detecting signatures of recent positive selection and background selection using pairwise coalescence times along the human genome. We first analyzed 113,851 British individuals from the UK Biobank data set^29^, detecting 12 loci with unusually high density of very recent coalescence times as a result of recent positive selection at these sites. These include 6 known loci linked to nutrition, immune response, and pigmentation, as well as 6 novel loci involved in immunity, taste reception, and other aspects of human physiology. We then analyzed 498 unrelated WGS samples from the Genome of the Netherlands data set^30^ to search for signals of background selection at deeper time scales and finer genomic resolution. We determined that SNPs in regions with low values of average coalescence time are strongly enriched for heritability across 20 independent diseases and complex traits (average *N*=86k), even when conditioning on a broad set of functional annotations (including other background selection annotations).

## Results

### Overview of ASMC method

We developed a new method, ASMC, that estimates the coalescence time (which we also refer to as time to most recent common ancestor, TMRCA) for a pair of chromosomes at each site along the genome. ASMC utilizes a Hidden Markov Model (HMM), which is built using the coalescent with recombination process^7-11^; the hidden states of the HMM correspond to a discretized set of TMRCA intervals, the emissions of the HMM are the observed genotypes, and transitions between states correspond to changes in TMRCA along the genome due to historical recombination events. ASMC shares several key modeling components with previous coalescent-based HMM methods, such as the PSMC^1^, the MSMC^2^, and, in particular, the recently developed SMC++ ^3^. In contrast with these methods, however, ASMC’s main objective is not to reconstruct the demographic history of a set of analyzed samples. Instead, ASMC is optimized to efficiently compute coalescence times along the genome of pairs of individuals in modern data sets. To this end, the ASMC improves over current coalescent HMM approaches in two key ways. First, by modeling non-random ascertainment of genotyped variants, ASMC enables accurate processing of SNP array data, in addition to WGS data. Second, by introducing a new dynamic programming algorithm, it is orders of magnitude faster than other coalescent HMM approaches, which enables it to process large volumes of data. Details of the method are described in the **Online Methods** section; we have released open-source software implementing the method (see URLs).

### Simulations

We assessed ASMC’s accuracy in inferring locus-specific pairwise TMRCA from SNP array and WGS data via coalescent simulations using the ARGON software^31^. Briefly, we measured the correlation between true and inferred average TMRCA for all pairs of 300 individuals simulated using a European demographic model^3^, for a 30 Mb region with SNP density and allele frequencies matching those of the UK Biobank data set (**Figure 1**; see **Online Methods**). As expected, ASMC achieved high accuracy when applied to WGS data (*r*^2^=0.95). When sparser SNP array data were analyzed, the correlation remained high (e.g. *r*^2^=0.87 at UK Biobank SNP array density), and increased with genotyping density. Similar relative results were obtained when comparing the root mean squared error (RMSE) between true and inferred TMRCA at each site, and the posterior mean estimate of TMRCA attained higher accuracy than the maximum-a-posteriori (MAP) estimate (**Supplementary Figure 1**). Inferring locus-specific TMRCA is closely related to the task of detecting genomic regions that are identical-by-descent (IBD), i.e. regions for which the true TMRCA is lower than a specified cut-off; ASMC attained higher IBD detection accuracy (area under the precision-recall curve) than the widely used Beagle IBD detection method^32^ (**Supplementary Table 1**).

**Figure 1.**
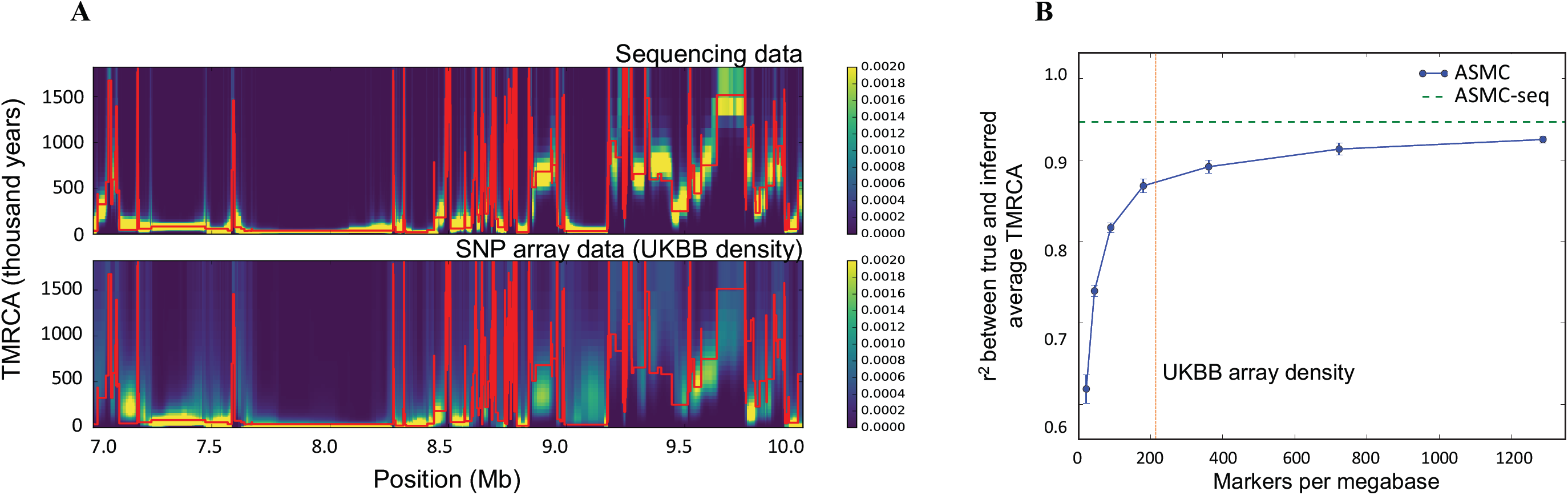
ASMC accuracy in coalescent simulations. (A) Sample posterior decoding of TMRCA along a 3 Mb segment for a pair of simulated individuals with ASMC run on WGS data (top) and on SNP array data (bottom). Red lines represent the true TMRCA, while the heat map represents the inferred posterior distribution. Posterior density tends to concentrate more tightly around the true TMRCA when WGS data are analyzed, due to the higher density of polymorphic variants. Posterior estimates using SNP array data are more dispersed for distant TMRCA, but remain highly concentrated for recent TMRCA. (B) Accuracy (*r*^2^ between true and inferred average TMRCA) as a function of marker density. TMRCA are inferred using the posterior mean obtained by ASMC at each site. ASMC-seq represents the accuracy obtained using ASMC on WGS data. The red vertical line indicates marker density in the UK Biobank data set. Errors bars represent standard errors. Numerical results are reported in **Supplementary Table11**

We evaluated the robustness of ASMC to various types of model misspecification, including an inaccurate demographic model, inaccurate recombination rate map, and violations of the assumption of frequency-based SNP ascertainment. To evaluate the impact of using an inaccurate demographic model, we simulated data under a European demographic history, but assumed a constant effective population size when inferring TMRCA (see **Online Methods**). As expected, this introduced biases, decreasing the accuracy of inferred TMRCA as measured by the RMSE, but had a negligible effect on the correlation between true and inferred TMRCA (**Supplementary Table 2**). An inaccurate demographic model is thus likely to result in biased TMRCA estimates, but has little effect on the relative ranking of TMRCA along the genome. Consistent with this observation, IBD detection remained accurate when an incorrect demographic model was used (**Supplementary Table 3**). We used a similar approach to evaluate the impact of using an inaccurate recombination rate map (see **Online Methods**), observing only negligible effects on the accuracy of inferred TMRCA (**Supplementary Table 4**). We next tested the robustness of ASMC to violations of the assumption that observed polymorphisms are ascertained solely based on their frequency, by instead ascertaining more rare variants in certain regions (mimicking genic regions; see **Online Methods**). We found that the distribution of inferred TMRCA in these “genic” regions did not deviate substantially from other regions (**Supplementary Figure 2**). Finally, we evaluated the impact of varying the number s of discrete TMRCA intervals (i.e. states of the HMM); we observed that increasing s had only a minor impact on posterior mean estimates of TMRCA, although the higher resolution led to noisier MAP estimates (**Supplementary Table 5**).

We next evaluated the running time and memory cost of ASMC. Letting s be the number of discrete TMRCA intervals (i.e. states of the HMM) and *m* be the number of observed polymorphic sites, ASMC has asymptotic running time O(*sm*). In comparison, the SMC++ method, which was shown to be more computationally efficient than other coalescent-based methods^3^, has asymptotic running time O(*s*^*3*^*m*). Accordingly, we observed that the running time of ASMC was 2 to 4 orders of magnitude faster than SMC++ when applied to simulated WGS data, depending on the number of discrete TMRCA intervals (**Figure 2**). For example, analysis of a pair of simulated genomes using 100 discrete time intervals required 7.4 seconds on a single processor for ASMC, compared to 3.3 hours for SMC++. This speedup does not involve a significant loss in accuracy (**Supplementary Figure 3**). The memory cost of ASMC was also efficient compared to SMC++, scaling linearly with s (**Supplementary Figure 4**).

**Figure 2.**
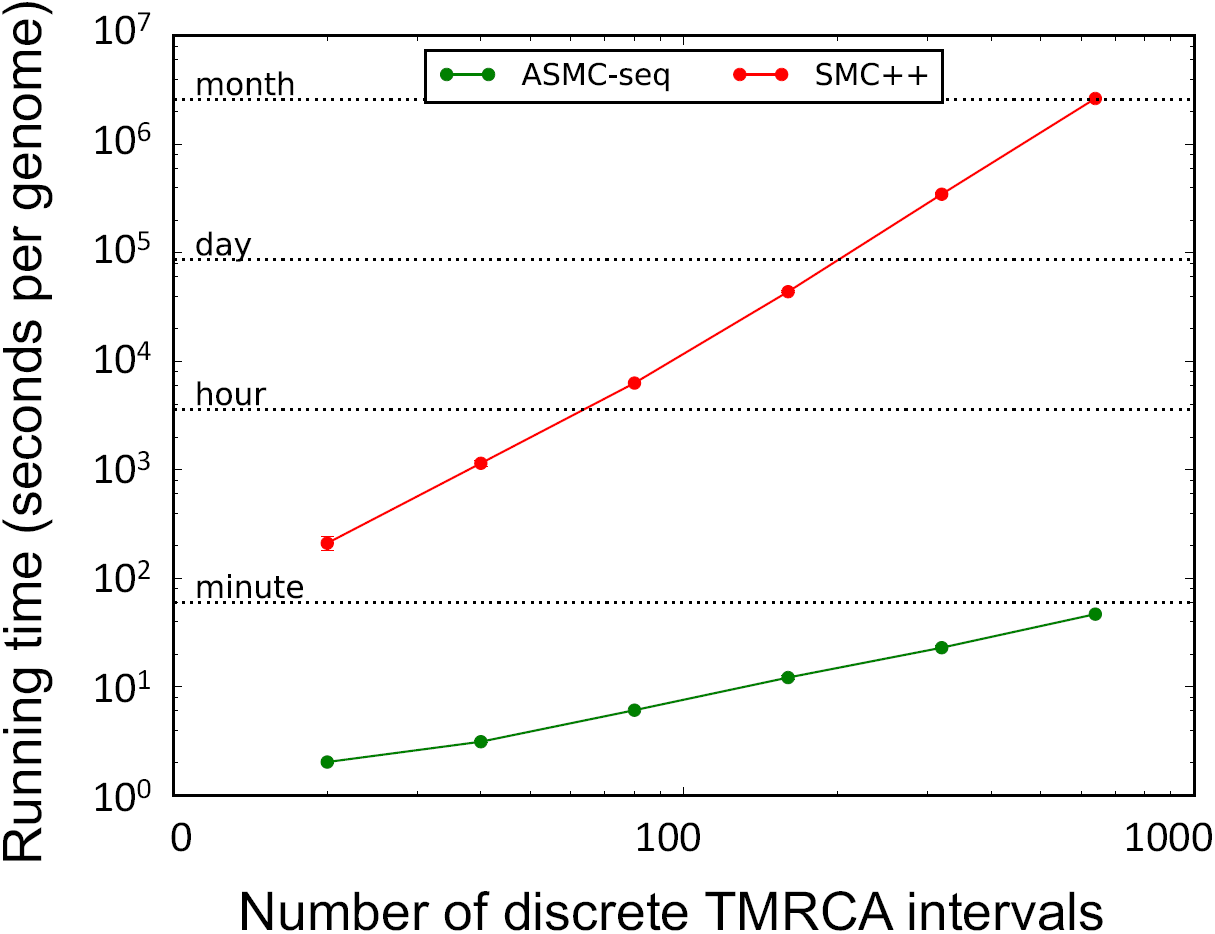
Running time of ASMC. We report the running time required to analyze a pair of simulated haploid genomes (extrapolated from running times in 5Mb regions) as a function of the number of discrete TMRCA intervals. Both SMC++ and ASMC-seq were run on WGS data. Numerical results are reported in **Supplementary Table 12**.

### Application to 113,732 samples from the UK Biobank reveals signals of recent positive selection

ASMC’s computational efficiency enables its application to analyses of TMRCA in large data sets. We thus used ASMC to infer locus-specific TMRCA in 113,732 unrelated individuals of British ancestry from the UK Biobank, typed at 678,956 SNPs after QC and phased using Eagle^33^ (see **Online Methods**); we note that phasing accuracy in this data set is very high, with average switch error rate on the order of 0.3% (one switch error every ∼10 cM^33^). We partitioned the data into batches of approximately 10,000 samples each and inferred locus-specific TMRCA for all haploid pairs within each batch, analyzing a total of 2.2 billion pairs of haploid genomes.

We sought to identify genomic regions with an unusually high density of very recent inferred TMRCA events (i.e. within the past several thousand years). Such signals are expected at sites undergoing recent positive selection, since a rapid rise in frequency of a beneficial allele causes all individuals with the beneficial allele to coalesce to a more recent common ancestor than under neutral expectation^34^; approaches to detect selection based on distortions in inferred coalescence times have been recently applied at different time scales^21^. We thus computed a statistic, DRC*T*, reflecting the *Density of Recent Coalescence* (within the past *T* generations), averaged within 0.05 cM windows. To compute approximate p-values, we noted that the DRC_*T*_ statistic under the null is approximately Gamma-distributed. We thus obtained approximate p-values for the DRC_*T*_ statistic by fitting a Gamma distribution to the null 18% of the genome obtained by conservatively excluding 500Kb windows around regions previously implicated in scans for positive selection (see **Online Methods**). Using coalescent simulations, we determined that DRC_150_ is highly sensitive in detecting signals of positive selection within the past ∼20,000 years, as compared to other methods^35,36^ (see **Online Methods, Supplementary Figure 5**).

Analyzing 63,103 windows of length 0.05cM in the UK Biobank data set, we detected 12 genome-wide significant loci (p < 0.05 / 63,103 = 7.9 × 10^−7^; see **Figure 3A and Table 1**). The loci that we detected exhibited strong enrichment of recent coalescent events spanning up to the past 20,000 years (**Figure 3B, 3C** and **Supplementary Figure 6**), consistent with our simulations (**Supplementary Figure 5**). Of the 12 loci, 6 are loci known to be under recent positive selection, harboring genes linked to nutrition (LCT^37^), immune response (HLA^38^, TLR^39^, IGHG^40^), eye color (GRM5^40^), and skin pigmentation (MC1R^40^). We also detected 6 novel loci, harboring genes involved in immune response (STAT4^41^, associated with autoimmune disease^42-44^); mucus production (MUC5B^45^ within cluster of mucin genes, involved in protection against infectious disease^43^, associated with several types of cancer^46^ and lung disease^47^); taste reception (PKD1L3^48^, associated with kidney disease^49,50^); cardiac and fetal muscle (MYL4, associated with atrial fibrillation^51^); blood coagulation (ANXA3^52^, associated with cancer^53^ and immune disease^54^); and brain-specific expression and immune response (FAM19A5^55^). We note that suggestive loci implicated by the DRC_150_ statistic (p < 10^−4^; **Supplementary Table 6**) include known targets of selection linked to eye color (HERC2^56,57^), retinal and cochlear function (PCDH15^40^), celiac disease (SLC22A4^57,58^) and skin pigmentation (SLC45A2^57^).

**Table 1.**
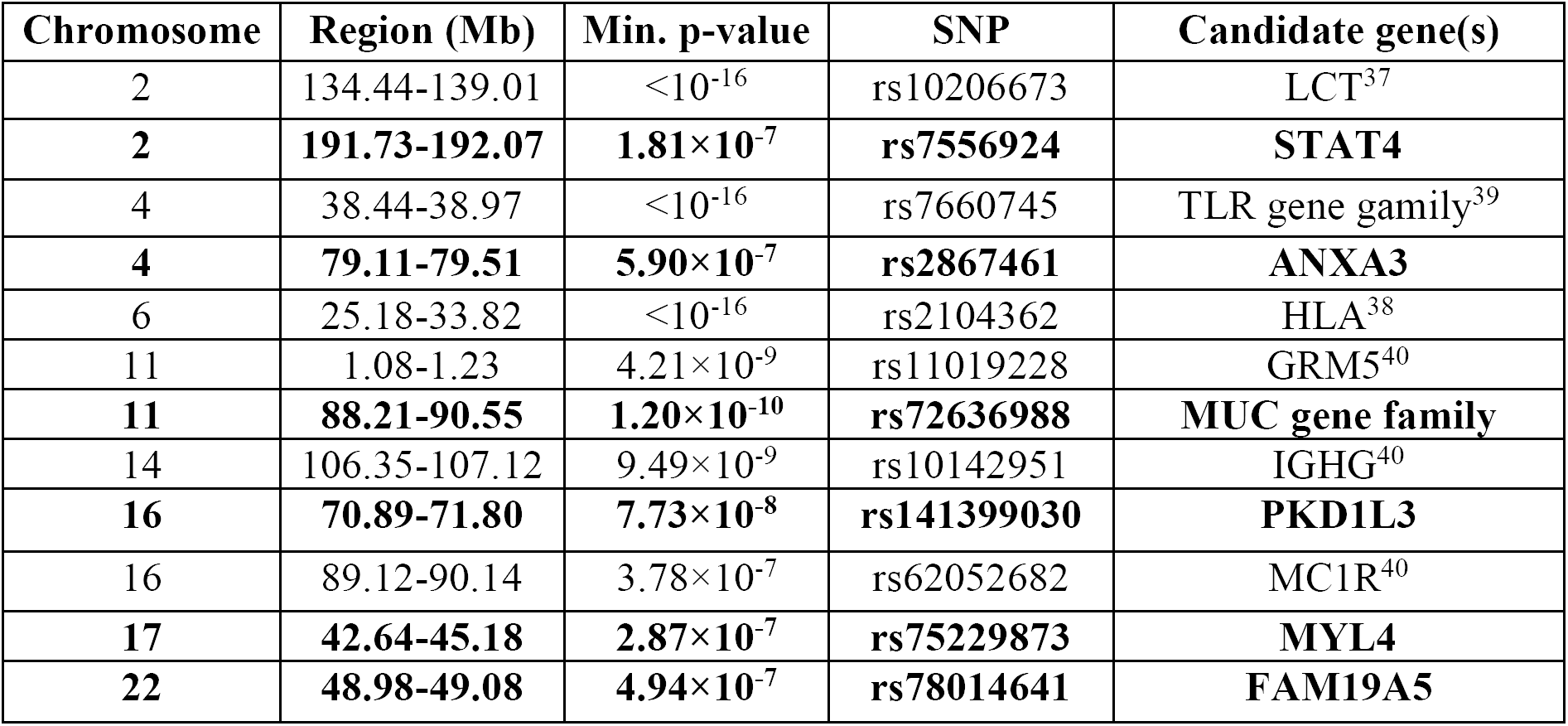
Genome-wide significant signals of recent positive selection. We report genomic locations, minimum p-value (capped at 10^−16^) across 0.05cM windows, SNP corresponding to signal peak, and candidate gene for the 12 genome-wide significant signals of recent positive selection (p < 0.05 / 63,103 = 7.9 × 10^−7^). Novel loci are denoted in bold font.

| Chromosome | Region (Mb) | Min. p-value | SNP | Candidate gene(s) |
| --- | --- | --- | --- | --- |
| 2 | 134.44-139.01 | $<10^{-16}$ | rs10206673 | LCT<sup>37</sup> |
| **2** | **191.73-192.07** | **$1.81 \times 10^{-7}$** | **rs7556924** | **STAT4** |
| 4 | 38.44-38.97 | $<10^{-16}$ | rs7660745 | TLR gene family<sup>39</sup> |
| **4** | **79.11-79.51** | **$5.90 \times 10^{-7}$** | **rs2867461** | **ANXA3** |
| 6 | 25.18-33.82 | $<10^{-16}$ | rs2104362 | HLA<sup>38</sup> |
| 11 | 1.08-1.23 | $4.21 \times 10^{-9}$ | rs11019228 | GRM5<sup>40</sup> |
| **11** | **88.21-90.55** | **$1.20 \times 10^{-10}$** | **rs72636988** | **MUC gene family** |
| 14 | 106.35-107.12 | $9.49 \times 10^{-9}$ | rs10142951 | IGHG<sup>40</sup> |
| **16** | **70.89-71.80** | **$7.73 \times 10^{-8}$** | **rs141399030** | **PKD1L3** |
| 16 | 89.12-90.14 | $3.78 \times 10^{-7}$ | rs62052682 | MC1R<sup>40</sup> |
| **17** | **42.64-45.18** | **$2.87 \times 10^{-7}$** | **rs75229873** | **MYL4** |
| **22** | **48.98-49.08** | **$4.94 \times 10^{-7}$** | **rs78014641** | **FAM19A5** |

**Figure 3.**
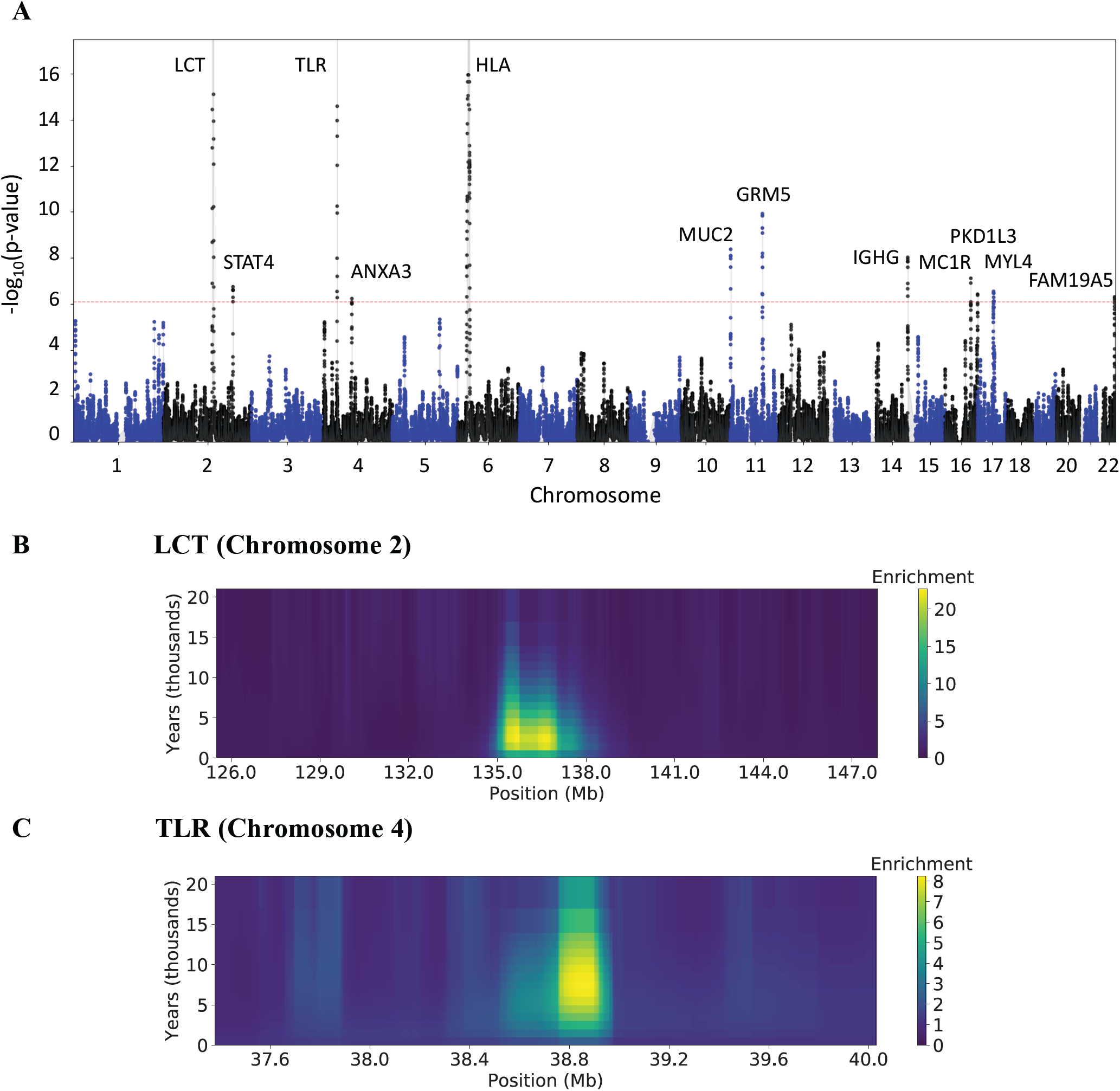
Genome-wide scan for recent positive selection in the UK Biobank data set. (A) Manhattan plot with candidate gene labels for 12 loci detected at genome-wide significance (p < 0.05 / 63,103 = 7.9 × 10^−7^; dashed red line). Numerical results for top loci are reported in Table 1; additional suggestive loci are reported in **Supplementary Table 6.** (B) Enrichment for recent coalescence events at the LCT locus (Chromosome 2). (C) Enrichment for recent coalescence events at the TLR locus (Chromosome 4). y-axis labels assume a 30-year generation time. Analogous plots for other top loci are provided in **Supplementary Figure 6**.

### Background selection annotation and heritability enrichment for complex traits

We next sought to detect signals of background selection at deeper time scales. To accomplish this, we used ASMC to estimate locus-specific TMRCA for all ∼0.5 million pairs of haploid genomes from unrelated individuals in the Genome of the Netherlands (GoNL) WGS data set (498 samples and 19,730,834 variants after QC; see **Online Methods**); we note that WGS data are required to achieve accurate resolution at deeper time scales (**Figure 1A**). Motivated by the fact that background selection reduces the effective population size at affected regions^34,59^, we estimated its strength by measuring the average pairwise TMRCA at each site, which is proportional to effective population size^60^. We refer to this background selection annotation as ASMC_avg_. The genome-wide average of ASMC_avg_ in the GoNL data was 17,399 generations (s.d. = 9,957 generations); we thus expect the ASMC_avg_ annotation to reflect background selection occurring within the past several hundred thousand years. As expected, ASMC_avg_ was highly correlated with other measures of background selection, including nucleotide diversity (r=0.50), the McVicker B-statistic^59^ (r=-0.28), and allele age predicted by ARGWeaver^6^, quantile-normalized within 10 minor allele frequency bins^61^ (r=0.26, see **Supplementary Table 7**).

Analyses using stratified LD score regression (S-LDSC)^62^ have shown that regions under background selection are enriched for disease and complex trait heritability^61^; enrichment was observed for the nucleotide diversity, McVicker B-statistic, and ARGWeaver predicted allele age annotations, as well as three other annotations linked to LD and recombination. We evaluated the ASMC_avg_ background selection annotation for heritability enrichment by applying S-LDSC to summary association statistics from 20 independent diseases and complex traits (**Supplementary Table 8**, average *N*=86k). We performed both an unconditioned analysis using only the ASMC_avg_ annotation, and a joint analysis conditioned on the 75 annotations from the baselineLD model^61^ (which includes a broad set of functional annotations, in addition to the six annotations linked to background selection and LD), in order to specifically assess whether our annotation provides additional signal. Focusing on the ASMC_avg_ annotation, we computed the *τ*\* metric^61^, defined as the proportionate change in per-SNP heritability resulting from a 1 standard deviation increase in the value of the annotation, conditional on other annotations included in the model.

In the unconditioned analysis, lower ASMC_avg_ was associated with higher per-SNP heritability for all 20 traits analyzed (**Figure 4A**), confirming that regions under background selection are enriched for disease heritability. Meta-analyzed across the 20 traits, the τ;* for ASMC_avg_ had a value of −0.81 (s.e. = 0.01; Z-test p < 10^−300^). After conditioning on the baselineLD model, the *τ*\* for ASMC_avg_ remained strongly significant at −0.25 (s.e. = 0.01; Z-test p = 7×10’^153^), implying that ASMC_avg_ remains informative for disease heritability after conditioning on other annotations linked to background selection as well as a broad set of functional annotations. Furthermore, ASMC_avg_ attained a larger value of *τ*\* than each of the other six annotations linked to background selection (**Figure 4B**), implying that it was the most disease-informative background selection annotation in this analysis; we note that adding ASMC_avg_ to the baselineLD model reduced the |*τ*\*| of the nucleotide diversity annotation from 0.13 to 0.00 and reduced the |*τ*\*| of the ARGWeaver^6^ predicted allele age annotation from 0.25 to 0.13, indicating that ASMC_avg_ subsumes signals from these annotations. We computed the proportion of heritability explained by each quintile of the ASMC_avg_ annotation, which provides a more intuitive interpretation of the strength of the annotation’s effect (**Figure 4C**). We observed that SNPs in the smallest quintile of the annotation explained 33.1% (s.e. 0.5%) of heritability, 3.8x more than SNPs in the highest quintile (8.7%, s.e. 0.5%), the largest ratio among annotations linked to background selection (**Supplementary Table 9**) (tied with the nucleotide diversity annotation, whose effect was however subsumed by the ASMC_avg_ annotation; Figure 4B). Annotations constructed based on average pairwise TMRCA conditional on the allele present on each chromosome were further informative for disease heritability (**Supplementary Figure 7** and **Supplementary Figure 8**; see **Online Methods**).

**Figure 4.**
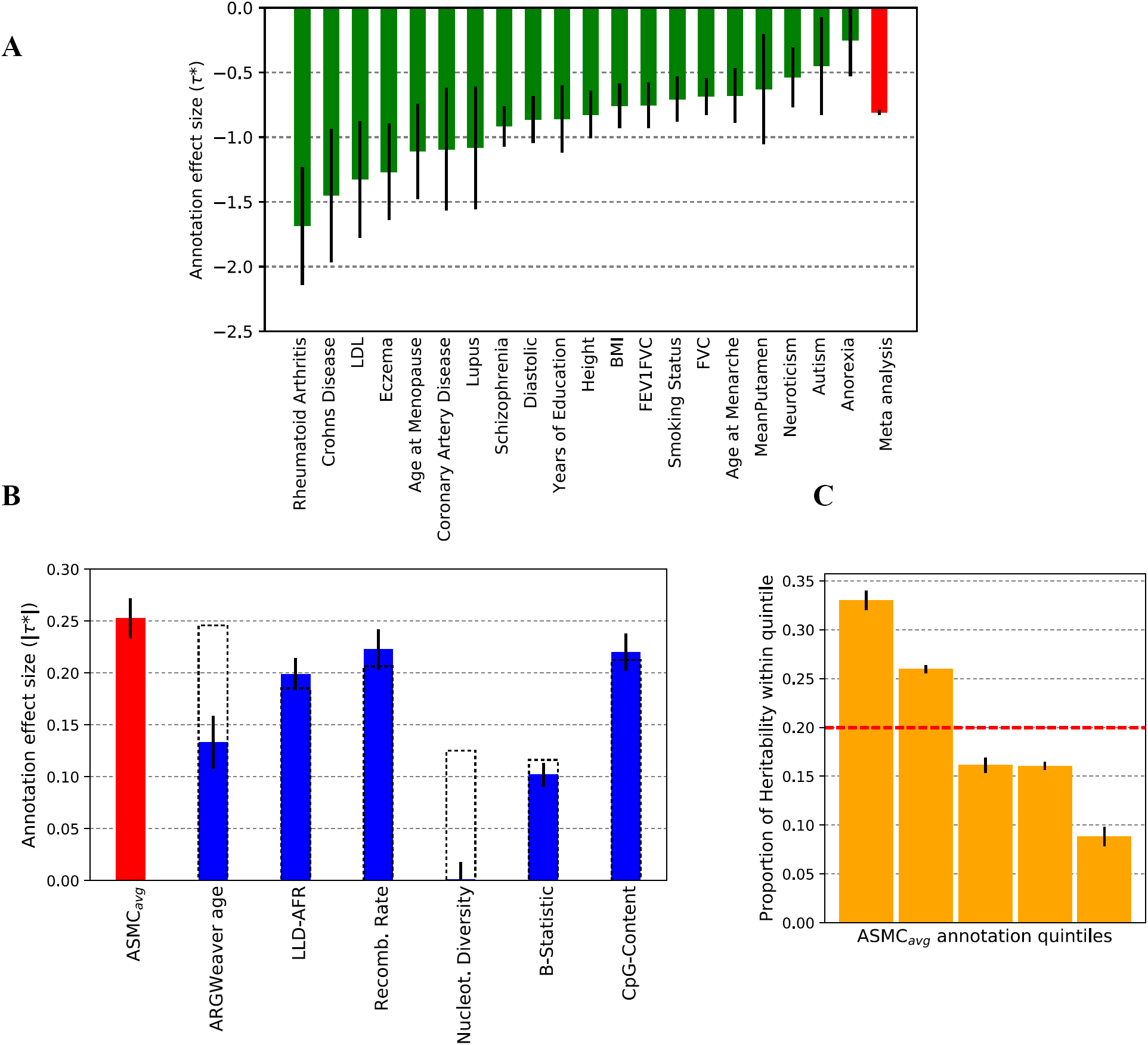
S-LDSC analysis of ASMC_avg_ background selection annotation and disease heritability. (A) *τ*\* value of the ASMC_avg_ annotation for 20 independent diseases and complex traits. (B) Absolute values of *τ*\* values (meta-analyzed across 20 independent diseases and complex traits) in joint analysis conditioned on baselineLD annotations. Dashed bars reflect values for six baselineLD annotations linked to background selection before the introduction of the ASMC_avg_ annotation. (C) Proportion of heritability explained by SNPs within different quintiles of ASMC_avg,_ annotation (in joint analysis conditioned on baselineLD annotations). Error bars represent standard errors. Numerical results are reported in **Supplementary Table 13**.

## Discussion

We have introduced a new method for inferring pairwise coalescence times, ASMC, that can be applied to either SNP array or WGS data and is highly computationally efficient. Exploiting ASMC’s speed, we analyzed ∼2.2 billion pairs of haploid chromosomes from 113,851 British samples within the UK Biobank data set, and detected strong evidence of recent positive selection at 6 known loci and 6 novel loci linked to immune response and other biological functions. We further used ASMC to detect background selection at deeper time scales, estimating the average TMRCA at each position along the genome of 498 WGS phased samples from the Netherlands. Using this annotation in a stratified LD score regression analysis of 20 diseases and complex traits, we detected a strong enrichment for heritability in regions predicted to be under background selection; our annotation had the largest effect among available annotations quantifying background selection.

High-throughput inference of ancestral relationships has a number of applications beyond those related to recent positive selection and disease heritability that we have pursued in this work. Genotype calling and imputation methods^25-28^, for instance, infer unobserved genotypes relying on ancestral relationships, which are usually estimated using computationally efficient approximations of the coalescent model (e.g. the *copying model*^63^). Related ideas have been applied to detect phenotypic associations^22-24^. The processing speed achieved by the ASMC approach, on the other hand, enables making minimal simplifications to the full coalescent process, while retaining high computational scalability. In addition, accurate detection of very recent common ancestors (IBD regions) across samples is a key component of several other types of analysis, including long-range phasing^33,64,65^, estimation of recombination rates using haplotype boundaries^66-68^, haplotype-based association studies^69^, estimation of mutation and gene conversion rates^70^, and inference of fine-scale demographic history within the past tens of generations^71-73^.

Although the ASMC offers new opportunities for inference of pairwise coalescence times, we note several limitations. First, the ASMC can operate either on pairs of unphased chromosomes within a single diploid individual, or on pairs of phased chromosomes across individuals. The latter application requires the availability of phased haplotypes, which may not be easy to obtain. We note, however, that recently developed methods for computational phasing attain sufficiently low switch error rates as to be considered negligible in most applications^33,64,74^. Second, ASMC assumes a demographic model that includes a single panmictic population, and does not allow for the presence of samples from multiple ethnic backgrounds. Analyses of multi-ethnic samples will require extending the current approach so that it can accommodate demographic models involving multiple populations. Third, ASMC is not currently applicable to imputed data. We have shown that higher genotyping density leads to higher accuracy in the inference of coalescence times. It may be possible to extend ASMC to incorporate information on imputation accuracy, enabling its application to imputed data. Fourth, our approach to assess the statistical significance of loci under recent positive selection is based on approximate p-value calculations. The use of approximate p-values has previously been adopted in detecting signals of positive selection^36^, and is more conservative than the widespread approach of simply ranking top loci^35^; nonetheless, the construction of an improved null model is a desirable direction of future development^75^. Finally, we note that although ASMC’s speed enables the analysis of large data sets, the computational costs of inferring pairwise coalescence times scale quadratically with the number of analyzed individuals. It may be possible to improve on this quadratic scaling given that at each location in the genome the ancestral relationships of a set of *n* samples can be efficiently represented using a tree-shaped genealogy containing n-1 nodes. The task of efficiently reconstructing a samples’ ancestral recombination graph (ARG)^6,24,76^, however, is substantially more complex than that of estimating pairwise TMRCA, and remains an exciting direction of future research. Despite these limitations and avenues for further improvement, we expect that ASMC will be a valuable tool for computationally efficient inference of pairwise coalescence times using SNP array or WGS data.

## Online Methods

We provide an overview of the main components of the ASMC approach. An extended description can be found in the **Supplementary Note**.

### ASMC model overview

The ASMC is a coalescent-based HMM^1-3,5^ (see **Supplementary Note** for background on related methods). At each site along the genome, hidden states represent the time at which a pair of analyzed haploid individuals coalesce, which we also refer to as their time to most recent common ancestor (TMRCA). In this model, time is discretized using a set of *s* user-specified time intervals, each representing a possible hidden state. The TMRCA may change between adjacent sites whenever a recombination event occurs along the lineages connecting the two individuals to their MRCA. The transition probability between states is modeled using a Markovian approximation^11^ of the full coalescent process. Observations are obtained using genotypes of the pair of analyzed samples, as well as a set of additional samples, as detailed below, and emission probabilities reflect the chance of observing a specific genotypic configuration, conditional on the pair’s TMRCA at a site. Calculations of the initial state distribution, the transition, and the emission probabilities consider the demographic history of the analyzed sample, which is separately estimated (e.g. using other coalescent HMMs run on available WGS data for the analyzed population) and provided as input. The main goal of the ASMC is to perform high-throughput inference of posterior TMRCA probabilities along the genome for many pairs of haploid individuals genotyped using either WGS or SNP array platforms.

### Emission model

ASMC’s emission probability calculations rely on the recently developed Conditioned Sample Frequency Spectrum (CSFS)^3^, which is extended to handle non-randomly ascertained genotype observations (e.g. SNP array data). Consider a sample of *n* individuals, and define 2 of them as *distinguished*, (*n*-2) of them as *undistinguished*. We are interested in estimating posterior TMRCA probabilities at a set of observed sites in the genome of the 2 distinguished samples. At each site, the CSFS model^3^ allows computing the HMM emission probability P(*d,u*|τ), i.e. the probability that *d*∈{0,1,2} derived alleles are carried by the distinguished pair of samples, while *u* ∈[0, n-2] derived alleles are observed in the (n-2) undistinguished samples, conditioned on the fact that the distinguished pair’s TMRCA (the HMM’s hidden state) is τ. Intuitively, this approach enables exploiting the relationship between an allele’s frequency and its age, which is modeled using the set of undistinguished samples and used to improve the inference of TMRCA for the distinguished pairs^3^. Because the set of undistinguished samples is solely used to obtain allele frequencies, their ancestral relationships need not be tracked, leading to a substantially simplified and tractable model. In the ASMC, this approach is extended to accommodate the fact that the observed sites may not be a randomly ascertained subset of polymorphic variants in the sample. To this end, we write the emission probability as P(obs|*d+u*)x*P*(*d,u|τ*), where the additional term P(obs|*d+u*) represents the probability that a site with (*d+u*) ∈[0, *n*] carriers of the derived allele is observed in the ascertained data. In the ASMC, this probability is estimated using the ratio between the empirical allele frequency spectrum obtained from the analyzed data and the allele frequency spectrum that is expected under neutrality for the demographic model provided in input. Details are provided in the **Supplementary Note**. The emission model enables handling both major/minor and ancestral/derived genotype data encoding. We verified using coalescent simulation (see **Simulations**), that the number of individuals used when computing the CSFS model does not have a substantial impact on accuracy (**Supplementary Table 10**).

### Transition model

The transition model describes the probability of transitioning along the genome between any pair of the s possible time intervals for the TMRCA of the two analyzed samples (which we referred to as *distinguished* individuals in the emission model). These transition probabilities are computed using the conditional Simonsen-Churchill model (CSC)^11,77^. In contrast to previously proposed Markovian approximations of the coalescent process, such as the SMC^9^ and the SMC’^10^, the CSC model remains accurate even if the observed genotypes are distant from one another^11^. This is an important requirement in the analysis of SNP array data, as markers in this type of data are separated by substantially larger genetic distances than in the case of WGS data. Details on the calculation of transition probabilities can be found in the **Supplementary Note**. ASMC supports variable recombination rates along the genome through a genetic map provided in input.

### Inference

The standard HMM forward-backward algorithm to perform posterior inference has computational cost O(s^2^m) for analysis using s hidden states in a sequence of length *m*^78^. Current analyses making use of coalescent HMMs to infer demographic histories utilize a number of hidden states in the order of 10^2^. When human WGS data is analyzed, the number of observed sites is in the order of 10^9^. Thus, the computational cost of applying the standard HMM approach is very high, and a number of solutions to speed up the inference have been proposed (see **Supplementary Note** for an overview). Here, we devise a new approach that uses dynamic programming to reduce the computational dependence on the number s of hidden states from quadratic to linear, resulting in a gain of 2 orders of magnitude for the average analysis compared to the standard algorithm. A related procedure exists for the SMC transition model^79^, but cannot be applied to the more accurate and more complex CSC approach used in this work. The speed-up in the HMM forward algorithm is obtained by simplifying the key operation of computing an updated *α’* vector of forward probabilities using the current forward vector, *α*, and the transition matrix, T, which is obtained from the CDC model. Computing the *i*-th entry of this vector normally requires performing the summation 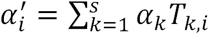, which has computational cost O(*s*). This operation, however, can be rewritten as a linear combination of three terms, each of which can be recursively computed in time O(1), reducing the cost of computing the entire forward vector from O(s^2^) to O(*s*) (see the **Supplementary Note** for a detailed derivation). An equivalent speed-up can be obtained for the backward algorithm. Furthermore, to reduce the dependence of ASMC’s running time on sequence length when WGS data are analyzed, we make the following approximation. Consider two polymorphic sites separated by a stretch of *n* monomorphic sites. Computing an updated forward probability vector *α’* using the standard approach would require performing the operation *α’* = α(TE_0_)^n^TEp, where E_0_ is a diagonal matrix with emission probabilities for a monomorphic site in its diagonal entries and Ep is the equivalent matrix for the emission at the next polymorphic site in the sequence. For short genetic distances that are observed between polymorphic sites, the matrix T is close to diagonal, and we can thus effectively approximate this product as *α*T^n^E^n^_0_TEp (see **Supplementary Figure 3).** Using the previously described dynamic programming approach, this operation can be computed in time O(*s*), and only needs to be performed at a subset of polymorphic sites, resulting in a further speedup of 2-3 orders of magnitude compared to the standard forward/backward approach operating on all sites. This approximation is not needed when SNP array data are analyzed, as we need not integrate over large stretches of monomorphic sites, treating instead all sites between a pair of genotyped SNPs as unobserved. In addition to this, most quantities involved in the O(*ms*) forward/backward operations can be precomputed and stored in a cache, substantially reducing constant terms in the computation.

### ASMC simulations

We performed extensive coalescent simulations to assess the accuracy of the ASMC method. Unless otherwise specified, all simulations use the setup described in this section (standard setup). We used the ARGON simulator^31^, incorporating a human recombination rate map (see **URL**s) and a recent demographic model for European individuals^3^. We simulated 300 haploid individuals and a region of 30Mb. To simulate SNP array data, we subsampled polymorphic variants to match the genotype density and allele frequency spectrum observed in the UK Biobank data set (described below). We used recombination rates from the first 30Mb of Chromosome 2, whose average rate of 1.66 cM/Mb well represents the recombination rates observed along the genome (mean 1.45 cM/Mb, s.d. 0.33 cM/Mb across autosomes). The demographic model and genetic map used to simulate the data were used when running ASMC, unless otherwise specified.

### Time discretization

We ran ASMC using different numbers of discrete time intervals, which were chosen to correspond to quantiles of the pairwise coalescence distribution induced by the demographic model. To achieve increased resolution into the recent past, some simulations utilized 160 discretization intervals chosen as follows: 40 intervals of length 10 between generations 0 and 400, 80 intervals of length 20 between generations 400 and 2,000, and 40 intervals corresponding to quantiles of the coalescence distribution, starting at generation 2,000. While using a larger number of time intervals provides increased resolution, the choice of time discretization should take into account that a larger number of time intervals typically results in noisier MAP estimates of TMRCA (see **Supplementary Table 5)**.

### Accuracy evaluation

ASMC’s inference accuracy was evaluated using two metrics. For a given region, and for all pairs of samples in a simulated data set, we computed the squared correlation (*r*^2^) between the true and inferred sum of TMRCA at each site within the region. This metric captures the accuracy of inferred genetic kinship, but is unchanged by potential scaling factors and possible systematic biases in the TMRCA estimates. We thus also measured the root mean square error (RMSE) between true and inferred TMRCA at individuals sites, which we usually report as a percent difference compared to analysis of WGS data for improved readability. For our analysis of IBD detection accuracy, we defined as true IBD regions all sites for which pairwise TMRCA were lower than a specified time threshold. We ran Beagle^32^ (v4.1) providing the true genetic map and using default parameters, and used threshold values for the output LOD score *(ibdlod)* to select the set of inferred IBD sites. To detect IBD using ASMC, we obtained MAP estimates of TMRCA at all sites using 160 discretization intervals (see **Time discretization**), and used thresholds on the inferred TMRCA values to select the set of inferred IBD sites. For both methods, we computed accuracy using the precision-recall curve. Neither Beagle nor ASMC enable obtaining recall values in the full [0,1] range, due to the presence of a lower bound for Beagle’s admissible LOD threshold values, and ASMC’s time discretization. To compare the two methods’ accuracies in each simulation, we computed the area under the precision-recall curve (auPRC) only within the range in which the accuracy of both methods could be measured, and reported the percent difference between the two methods’ auPRC (see **Supplementary Figure 9** for an illustration). The PRC curve between observed points was interpolated using the method of ref ^80^.

### Model misspecifications

To test ASMC’s robustness to an accurate demographic model we simulated data under a European demographic history, but ran ASMC assuming a constant population size of 10,000 diploid individuals (see **Supplementary Table 2**). To mimic inaccuracies in the genetic map we simulated data using a human recombination map for the simulated region, but run ASMC using a map with added noise. To introduce noise, the recombination rate between each pair of contiguous markers in the map was altered by randomly adding or subtracting a fraction of its true value (see **Supplementary Table 4**). To test whether deviations from the assumption of frequency-based ascertainment introduce significant biases, we mimicked over-ascertainment of rare variants in genic regions of the genome. To this end, we randomly sampled ∼25% of the markers from 10Kb-long genes placed every 200Kb, while the remaining variants were sampled to match the UK Biobank frequency spectrum as in standard simulations, and compared the distribution of coalescent times within over-ascertained regions and the rest of the genome (**Supplementary Figure 2**).

### UK Biobank (UKBB) data set

The UK Biobank interim release data comprise 152,729 samples, from which we extracted 113,851 individuals of British ancestry (as described in ref. ^81^). 95 trio parents were excluded and used to assess phasing quality with the Eagle^33^ software, leaving a total of 113,756 samples. From these, we created 11 batches with 10,000 samples and 1 batch with the remaining 3,756 samples, which we analyzed using ASMC. Out of the original ∼800k variants (for basic quality control details see **URLs**: UK Biobank Genotyping and QC), we analyzed a total of 678,956 SNPs that were autosomal, polymorphic in the set of analyzed samples, biallelic, with missingness ≤10%, and not included in a set of 65 variants with significantly different allele frequencies between the UK BiLEVE array and the UK Biobank array. We divided the genome in 39 autosomal regions from different chromosomes or separated by centromeres.

### Detection of recent positive selection

To detect the occurrence of recent positive selection, we computed a statistic related to the *Density of Recent Coalescence* events within the past *T* generations (DRC_*T*_ statistic). The DRC_*T*_ statistic was measured as follows. At a given site along the genome, we first averaged all posterior TMRCA estimates obtained from all analyzed pairs of samples and renormalized these averages to obtain an average pairwise coalescence distribution at the site. The DRC*T* statistic was then obtained by integrating this distribution between generations 0 and *T*. The statistic was measured in windows of 0.05 cM, reporting an average of all SNPs within each window.

### Null model calibration

Given *n* samples from a population of recent effective size *N*, the DRC_*T*_ statistic is approximately Gamma-distributed under the null for sufficiently small values of *T* and *n* ≪N. The rationale of this approximation is that for *n* ≪*N*, a small number of coalescence events will have occurred within the short time span of *T* generations. In this regime, the coalescence time of each pair of lineages may be modeled as independent and exponentially distributed, which allows approximating the total number of *early* coalescence events as a Gamma-distributed random variable. Similar approximations have been recently used elsewhere^36,82^. We thus computed approximate p-values for our selection scan in the UKBB data set using the following approach. We first extracted a subset of “neutral” genomic regions, spanning a total of 18% of the genome, and defined as any genotyped site at a distance greater than 500Kb from regions contained in a recent database of positive selection^83^ (see **URLs**: database of positive selection). We then built an empirical null model by fitting a Gamma distribution (using Python’s Scipy library, see **URLs**) to these putatively neutral regions, and used this model to obtain approximate p-values throughout the genome. We analyzed 63,103 windows, using a Bonferroni significance threshold of 0.05 / 63,103 = 7.9 × 10^−7^. One of the genome-wide significant signals that we detected (PKD1L3 locus, chr16:70.89-71.80Mb) fell within the putatively neutral portion of the genome. We thus iterated this procedure, excluding this locus from the set of putatively neutral loci.

### Selection simulations

We used the simulation setup recently adopted by Field et al.^36^ to test the sensitivity of the DRC*T* statistic in detecting recent positive selection, and its specificity to recent time scales. We simulated several replicates for a region of 10Mb and 6,000 haploid individuals from a European demographic model^19^, using the COSI2 coalescent simulator^84^. An allele at the center of the region was simulated to undergo recent positive selection, reaching a high present-day frequency of 0.7. We used the simuPOP^85^ software to obtain allele frequency trajectories under additive selection models, for several values of the selection coefficient. To test for specificity to recent time scales, we varied the period during which selection was active, posing no constraints on whether selection acted on a novel allele or on standing variation.

To assess power, we simulated 50 independent replicates for positive selection occurring in the past 200 generations (or ∼6,000 years), using selection coefficients S=0.01, 0.03, 0. 05, 0.1. We detected positive selection using either iHS (ref. ^35^), SDS (ref. ^36^), or DRC_150_ (**Supplementary Figure 5a**). The iHS statistic was computed using the Selscan software^86^ with default parameters. We computed the iHS statistic at either the sequenced causal variant (iHS_sequence_), or averaged at SNPs within a 0.05 cM window around the causal variant in simulated SNP array data (his_array_), which we obtained from simulated sequencing data as detailed above for neutral simulations. The DRC_150_ statistic was similarly computed by averaging within a 0.05 cM window on SNP array data. The SDS statistic was computed at the sequenced causal variant (SDS_sequence_). We found the DRC_150_ statistic computed on SNP array data to be highly sensitive to recent positive selection starting at S=0.03. Similar results for DRC_20_ are also reported in **Supplementary Figure 10a**.

To assess the specificity of DRC_150_ to recent time scale, we simulated selection starting at time -∞ and ending at a generation in 0, 50, 100, 200, 400, 600, 800, 1000, 1500, 2000 (**Supplementary Figure 5b**). We observed the DRC_150_ statistic to be mostly sensitive to selection acting during the past ∼700 generations (or ∼20,000 years), a similar time-span compared to the iHS statistic computed at the sequenced causal variant, which was however generally less sensitive, while the SDS statistic computed at the sequenced causal variant was only sensitive to extremely recent positive selection, as previously shown^36^. We also report DRC_20_ results in **Supplementary Figure 10b**.

We performed additional simulation to evaluate the calibration of the null model. We observed an excellent fit for the DRC_20_ statistic (**Supplementary Figure 11a**), and only moderate inflation for the DRC_150_ statistic (**Supplementary Figure 11b**). The amount of inflation observed in the empirical null model obtained using the DRC_150_ statistic within the UKBB data set was consistent with our coalescent simulations (**Supplementary Figure 11c,d**). We note that for very small values of *T* the independence assumption is more accurately met, so that the DRC*T* statistic is well approximated using a Normal distribution (see Supplementary Figure 12 for DRC_20_). We expect the moderate amount of inflation observed in neutral simulations for the DRC_150_ statistic to be counterbalanced in real data analysis by the conservative use of a Bonferroni significance threshold and the fitting of null model parameters using an empirical distribution of test statistics, which is likely to result in over-dispersion of the null model due to signals of positive selection that are too weak to be detected. Consistent with this hypothesis, genome-wide significant loci (Table 1) and suggestive loci (Supplementary Table 6) contain several regions of known recent adaptation.

### Genome of the Netherlands (GoNL) data set

The data set consists of 748 individuals who passed quality control and were sequenced at an average of ∼13x (quality control details for the Release 4 data are described elsewhere^30^). We analyzed 19,730,834 sequenced variants for 498 trio-phased unrelated parents, excluding centromeres and dividing the genome in the same 39 autosomal regions used for analysis of the UKBB data set.

### ASMC_avg_ annotation

We set out to estimate the strength of background selection by measuring variation in local effective population size along the genome^59^. We used ASMC to estimate the posterior mean TMRCA at all sites and for all pairs of haploid individuals in the GoNL data set. We averaged these estimates at each site to obtain an annotation of background selection, which we refer to as ASMC_avg_. We similarly computed other annotations, conditioning on whether either or both individuals at a site carried a mutated allele. The ASMC_het_ annotation (**Supplementary Figure 7**), was obtained by averaging at each site the posterior mean TMRCA estimates for all pairs of individuals that were found to be heterozygous at each site. Other annotations were similarly computed using only pairs carrying e.g. minor/major alleles at each site (see **Supplementary Figure 8**).

### Stratified LD Score (S-LDSC) analysis

We investigated whether large values of our annotations related to background selection corresponded to an enrichment in heritability for 20 complex traits and diseases listed in **Supplementary Table 8**. The S-LDSC analysis was run on data sets containing European individuals using standard guidelines^62^. The sets of LD-score, regression, and heritability SNPs were defined as follows. LD score SNPs were set to be 9,997,231 biallelic SNPs with at least 5 minor alleles observed in 489 European samples from the 1000 Genomes Phase 3 data set^12^ (see **URLs**); regression SNPs were set as 1,217,312 HapMap Project Phase 3 SNPs; and Heritability SNPs, used to compute trait heritability, were chosen as the 5,961,159 reference SNPs with MAF ≥ 0.05. The MHC region (2Mb 25-34 on Chromosome 6) and SNPs with, *χ*^2^>80 or 0.0001*N* were excluded from the analysis. Annotations contained in the baselineLD model, which we included in our joint analyses, can be found in Supplementary Table S9 of ref. ^61^. To avoid minor allele frequency (MAF)-mediated effects, all ASMC-related annotations used in the S-LDSC analysis were quantile-normalized with respect to MAF of regression SNPs. Specifically, we used 10 MAF ranges specified in the baselineLD model, corresponding to 10 frequency quantiles for the regression SNPs. For each range, we ranked values of an annotation for the corresponding SNPs, and mapped them to quantiles of a Standard Normal distribution. Annotation effects, *τ*\*, were obtained from the output of S-LDSC, as described in ref. ^61^. Independent traits were selected on the basis of low genetic correlation, as previously described^62^. Meta-analysis of *τ*\* values across independent traits was performed computing a weighted average of individual estimates of *τ*\*, weighted using 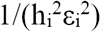 where 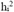 represents heritability for the *i*-th trait, and ε_*i*_ represents the standard error of the trait’s *τ*\* estimate.

## Acknowledgements

We thank Po-Ru Loh for suggesting several coding improvements for the ASMC software, and for support with the phasing and processing of the UK Biobank data; Steven Gazal for support with the S-LDSC analysis and the baselineLD model; Ilya Shlyakhter for support with the COSI2 simulator; Yair Field for support with the simulation setup in the analysis of recent positive selection; David Reich for providing computational resources; Hilary Finucane, Yakir Reshef, and Sasha Gusev for helpful discussions. This research was conducted using publicly available data sets (see URLs): the UK Biobank Resource under Application *#*16549, and the Genome of the Netherlands resource under Application *#*2017149. We thank the participants of the UK Biobank and the Genome of the Netherlands projects. P.P. and A.L.P. were supported by NIH grant R01 MH101244; J.T. and Y.S.S. were supported in part by an NIH grant R01-GM094402, and a Packard Fellowship for Science and Engineering; Y.S.S. is a Chan Zuckerberg Biohub investigator.

## URLs

- The ASMC program will be released prior to publication as a publicly available, open source software package at http://www.palamaralab.org/software/ and https://github.com/pierpal/ASMC
- UK Biobank website: http://www.ukbiobank.ac.uk/
- Genome of the Netherlands website: www.nlgenome.nl
- UK Biobank Genotyping and QC: http://www.ukbiobank.ac.uk/wp-content/uploads/2014/04/UKBiobank_genotyping_QC_documentation-web.pdf
- Human genetic maps: http://www.shapeit.fr/files/genetic_map_b37.tar.gz
- The dbPSHP database of positive selection: ftp://jjwanglab._org/db_PSHP/curation/dbP_SHP_20131001.tab
- Python’s Scipy library: http://www.scipy.org/
- 1000 Genomes Project Phase 3 data: ftp://ftp.1000genomes.ebi.ac.uk/vol1/ftp/release/20130502.

## Supplementary Figures and Tables

**Supplementary Figure 1.**
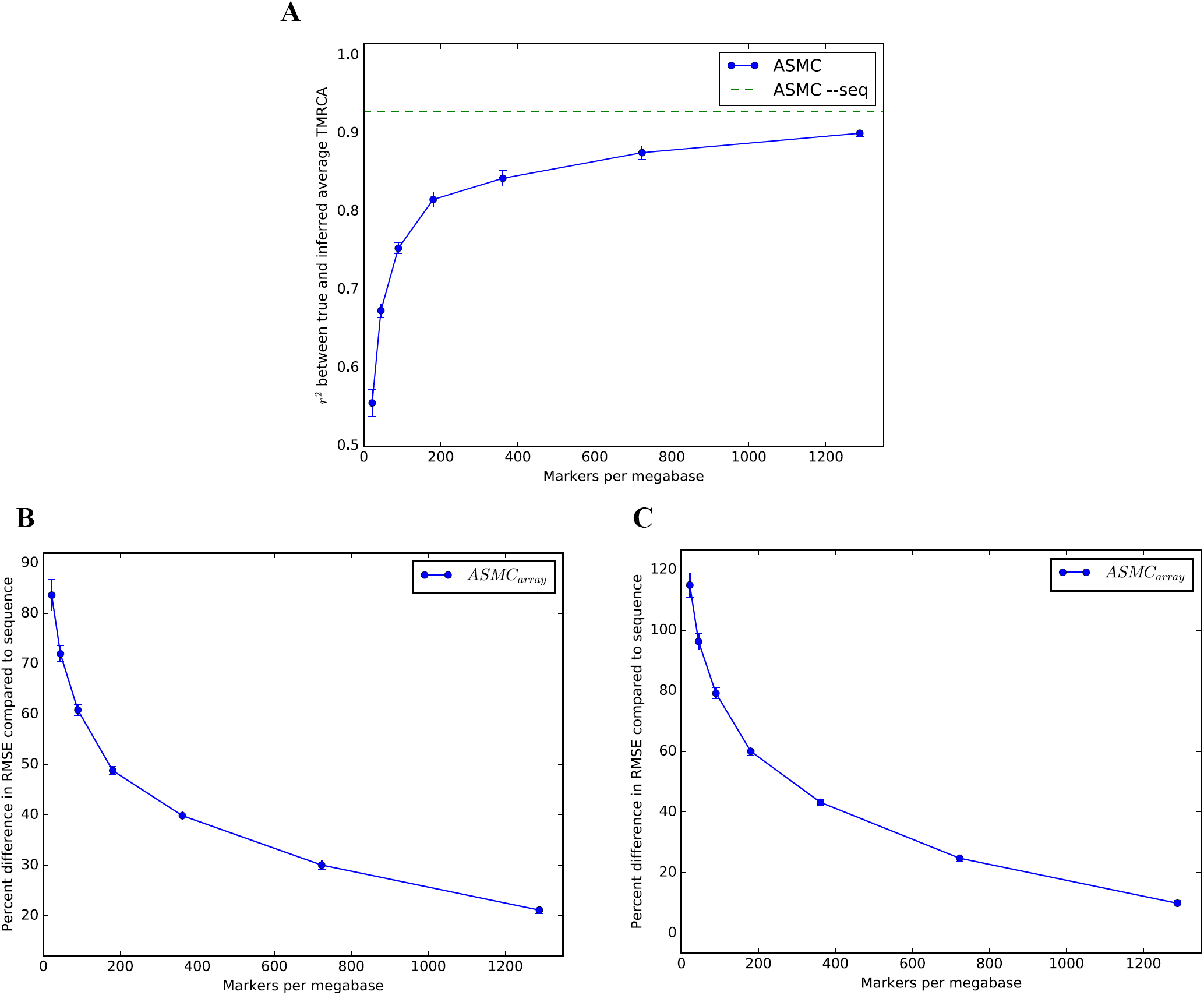
ASMC performance in simulations as a function of marker density. (A) *r*^2^ as a function of marker density, between true average TMRCA within the simulated region and average TMRCA inferred using the maximum-a-posteriori (MAP) of the posterior distribution computed by ASMC. ASMC-seq represents the accuracy obtained using ASMC on WGS data. Error bars represent standard errors. (B) We measure RMSE between true TMRCA at each site, and the TMRCA inferred by ASMC using the posterior mean on either SNP array or WGS data. We report the percent difference in per-site RMSE between analysis of SNP array data and WGS data (C) We measure RMSE between true TMRCA at each site, and the TMRCA inferred by ASMC using the MAP on either SNP array or WGS data. We report the percent difference in per-site RMSE between analysis of SNP array data and WGS data.

**Supplementary Figure 2.**
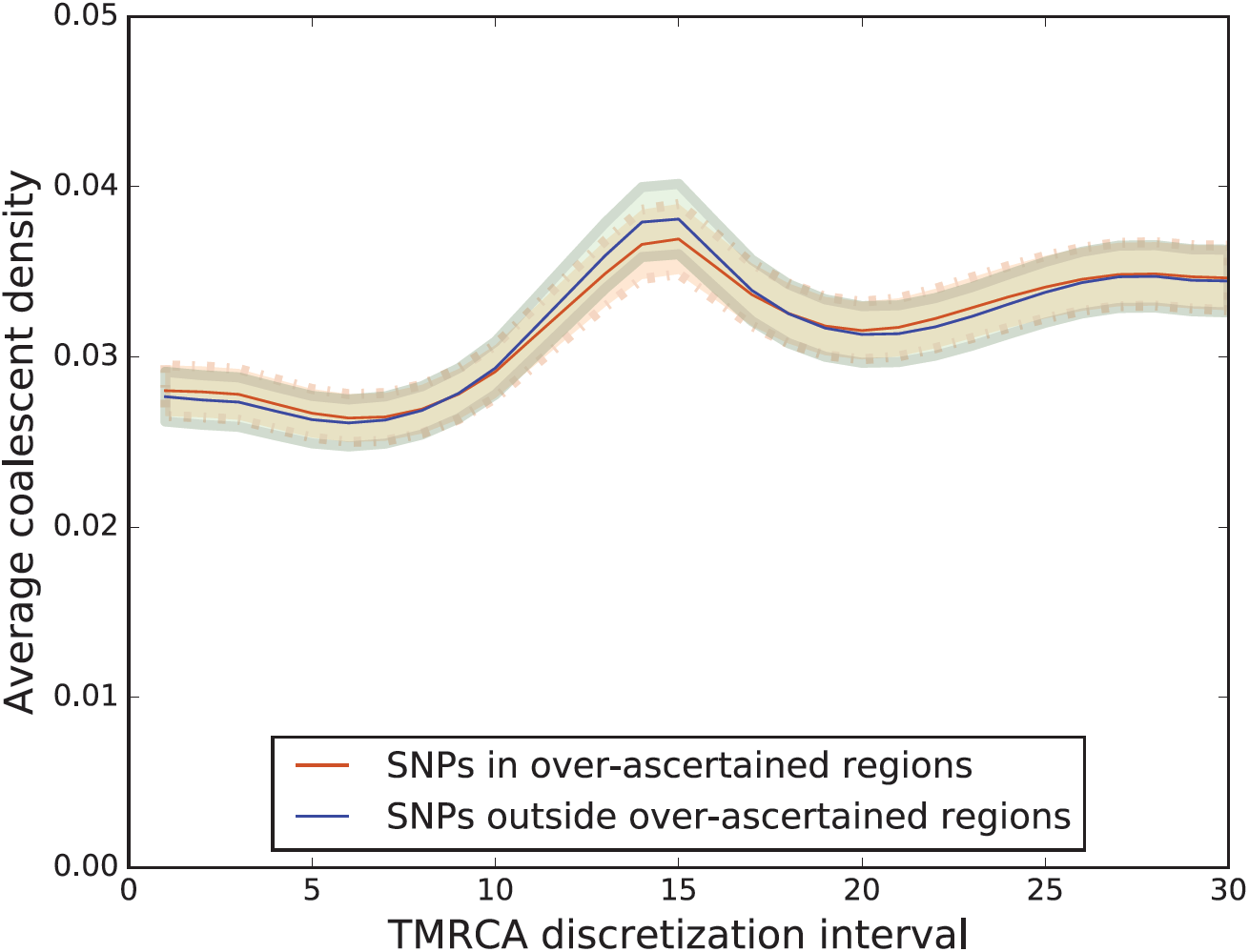
Robustness to deviations from frequency-based ascertainment. Approximately 25% of the variants found on the UK Biobank Axiom Array were selected based on their functional relevance, particularly in coding regions, while the remaining ∼75% were ascertained based on frequency. To mimic this ascertainment scheme in our simulations, we randomly sampled ∼25% of the markers from 10Kb-long genes placed every 200Kb, while the remaining variants were sampled to match the UK Biobank frequency spectrum as in standard simulations. Simulations were performed using the standard setup and 30 discretization intervals for TMRCA inference. Shaded regions indicate 95% confidence intervals. We observed minimal deviation between coalescence densities inferred within and outside the simulated gene regions

**Supplementary Figure 3.**
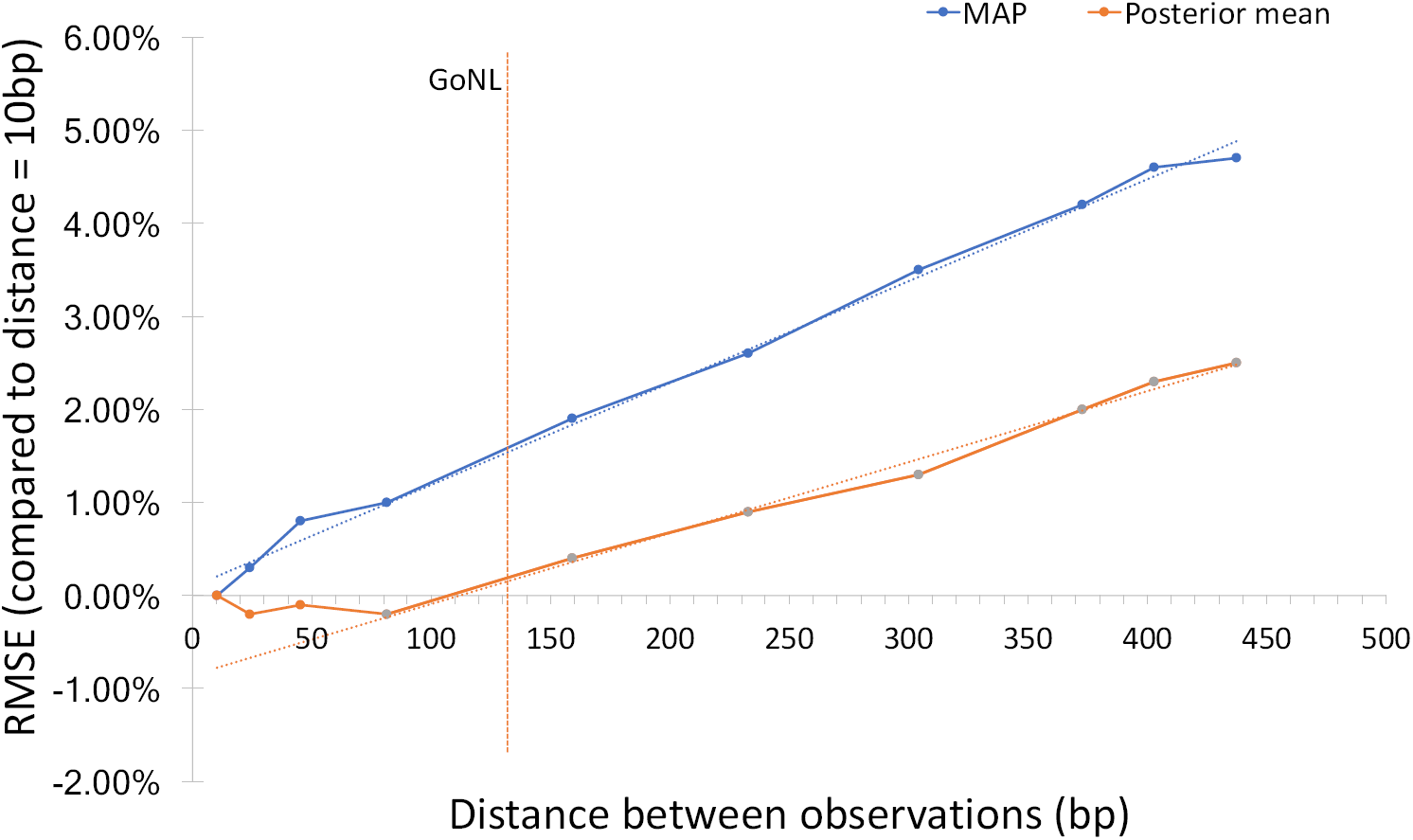
Effects of ASMC-seq transition approximation. When computing forward-backward probabilities in WGS data, ASMC-seq makes the approximation 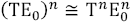, where *n* is the number of sites between two consecutive observations in the sequencing data, Eo is a diagonal matrix reflecting emission probabilities for *n* monomorphic sites, and T is the transition matrix between two sites at distance *n*, which is close to diagonal with off-diagonal entries growing with *n.* Exact calculations are obtained for *n* = 1 (or when ASMC is run on SNP array data), while an approximation is made for *n>* 1. To measure the extent to which this approximation affects inference accuracy, we measured the per-site RMSE between true TMRCA and TMRCA inferred using either maximum-a-posteriori (MAP) or posterior mean. We simulated 100 European samples in a 10 Mb region at the beginning of Chromosome 2 (with recombination rate 2.18 cM per Mb), and randomly inserted monomorphic sites along the genome to measure accuracy at different values of *n*, running ASMC-seq with 30 time intervals. We report RMSE for different values of *n*, as a percentage of the RMSE measured for *n* = 10, The red vertical bar represents the genotyping density observed for the GoNL data set *(n=* 136). For MAP inference, the error linearly increased at a rate of ∼0.01% per base pair, remaining below 2% for a genotyping density similar to the GoNL data set. For posterior mean inference, a negligible difference in accuracy was observed for *n<*100, followed by a linear increase at an approximate rate of ∼0.008% per base pair, and increased error below 0.5% at GoNL genotyping density.

**Supplementary Figure 4.**
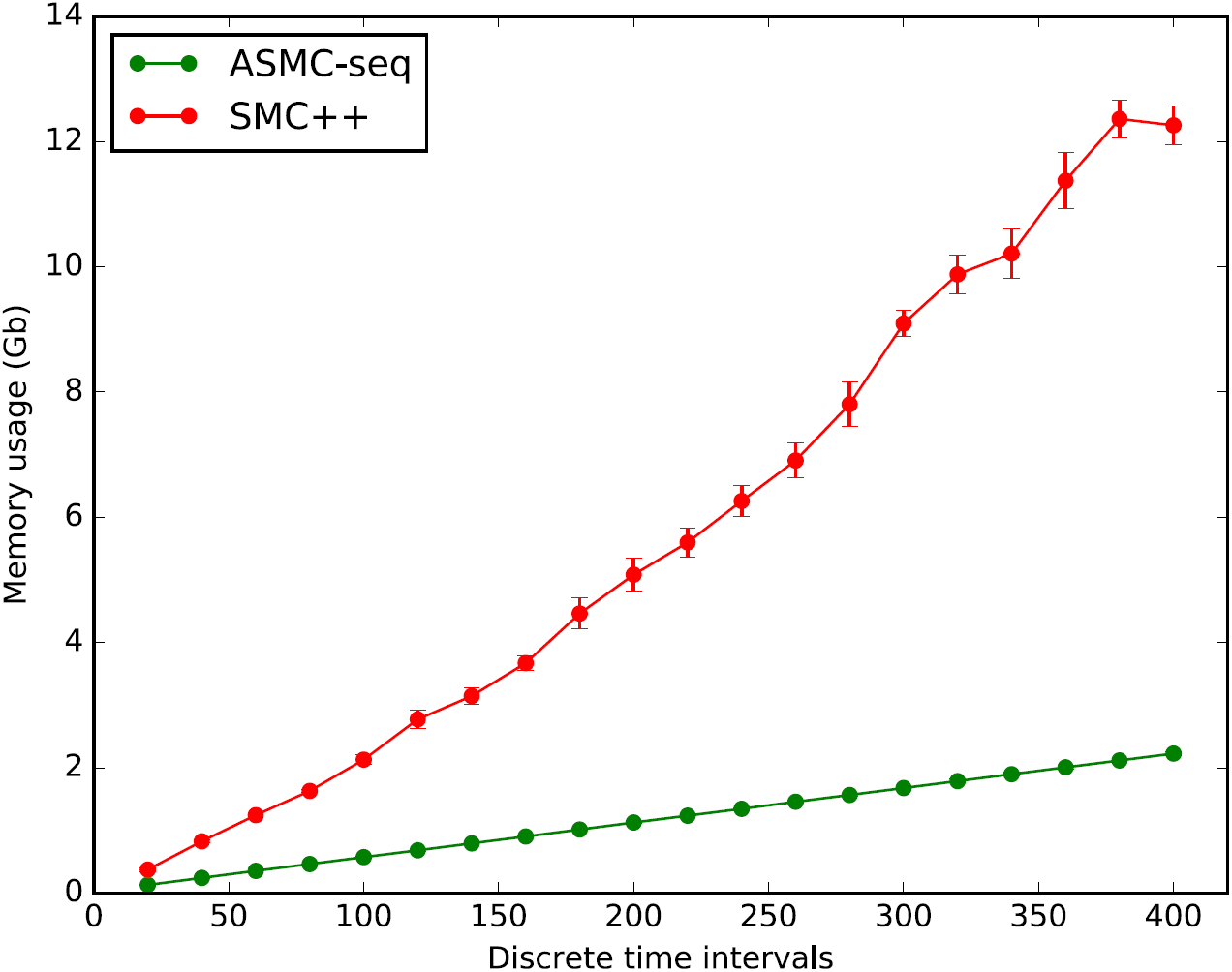
Memory use of ASMC-seq and SMC++. Memory usage for the analysis of coalescence times in a 5Mb region using WGS data from 100 haploid individuals. Bars represent standard errors.

**Supplementary Figure 5.**
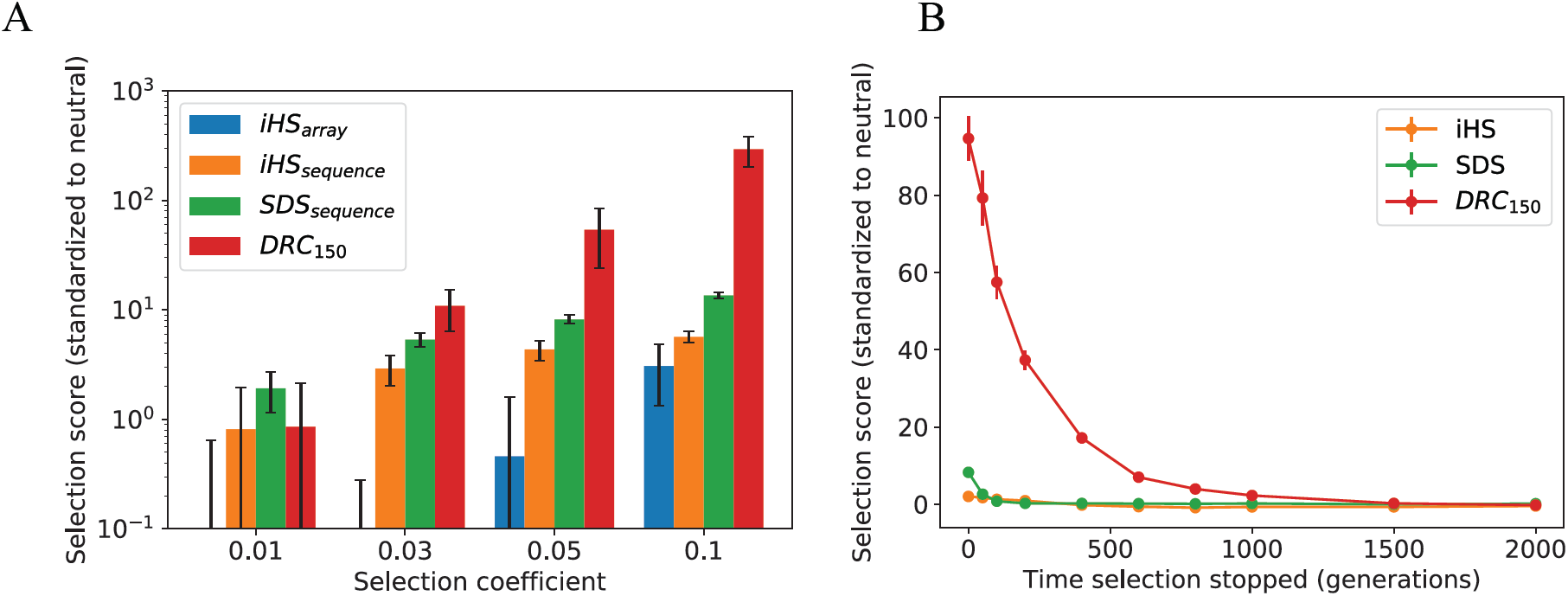
Selection simulations for DRC_150_. (A) Simulation of different strengths of recent positive selection starting 200 generations in the past: iHS score^35^ run on array data (iHSarray); iHS score^35^ on causal variant from sequencing data (iHS_sequence_); SDS score^36^ on causal variant from sequencing data (SDS_sequence_); DRC_150_ score on array data. Scores of each method are standardized with respect to corresponding scores obtained in neutral simulations. Bars indicate standard deviations. (B) Specificity to recent past for iHS and SDS run son sequencing data, and for DRC_150_. Simulation of selection starting at time -∞ stopping at the specified generation, followed by neutral drift. The DRC_150_ statistic is mostly sensitive to selection that has been active within the past ∼700 generations (or ∼20,000 years). Bars represent standard errors.

**Supplementary Figure 6.**
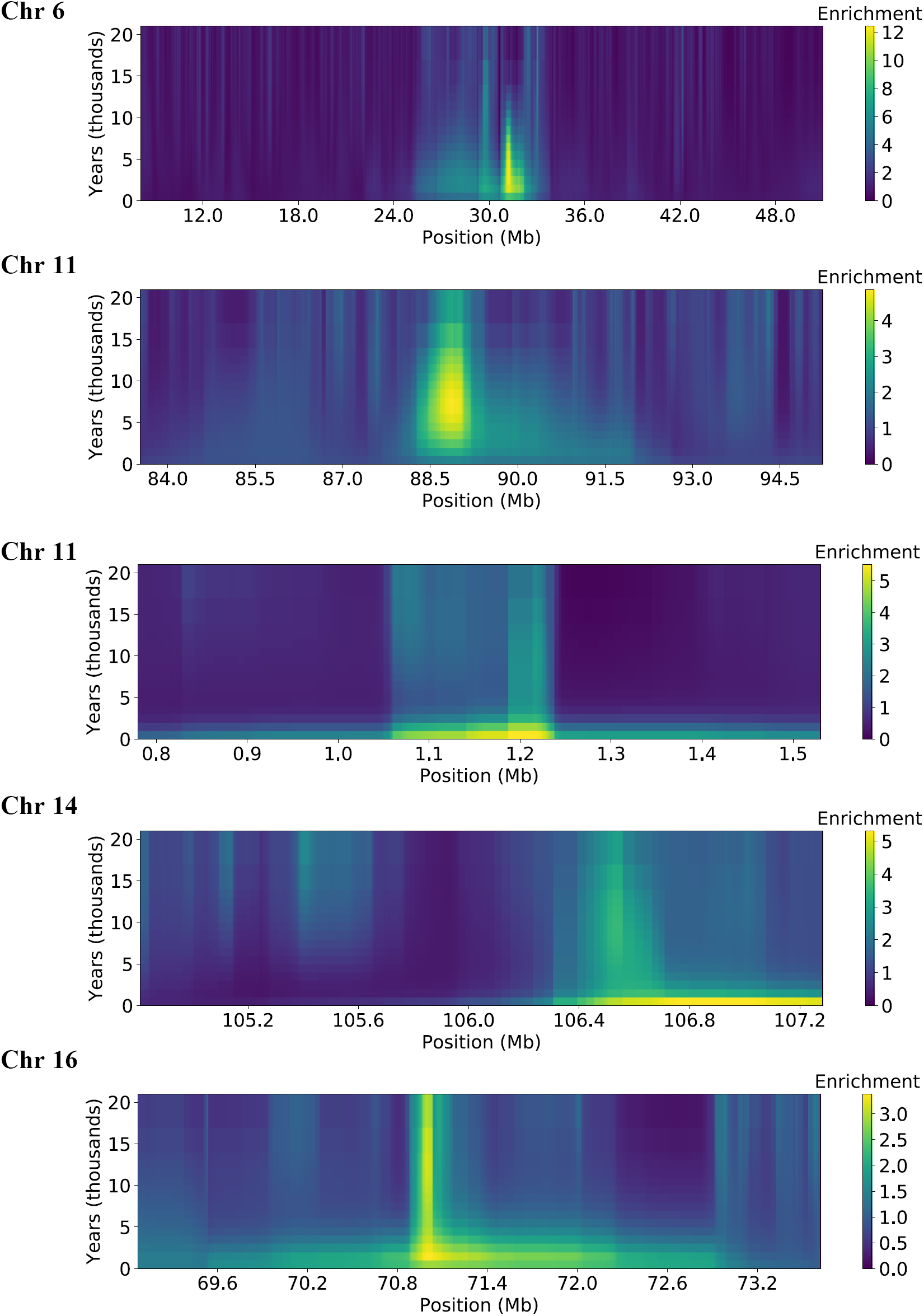
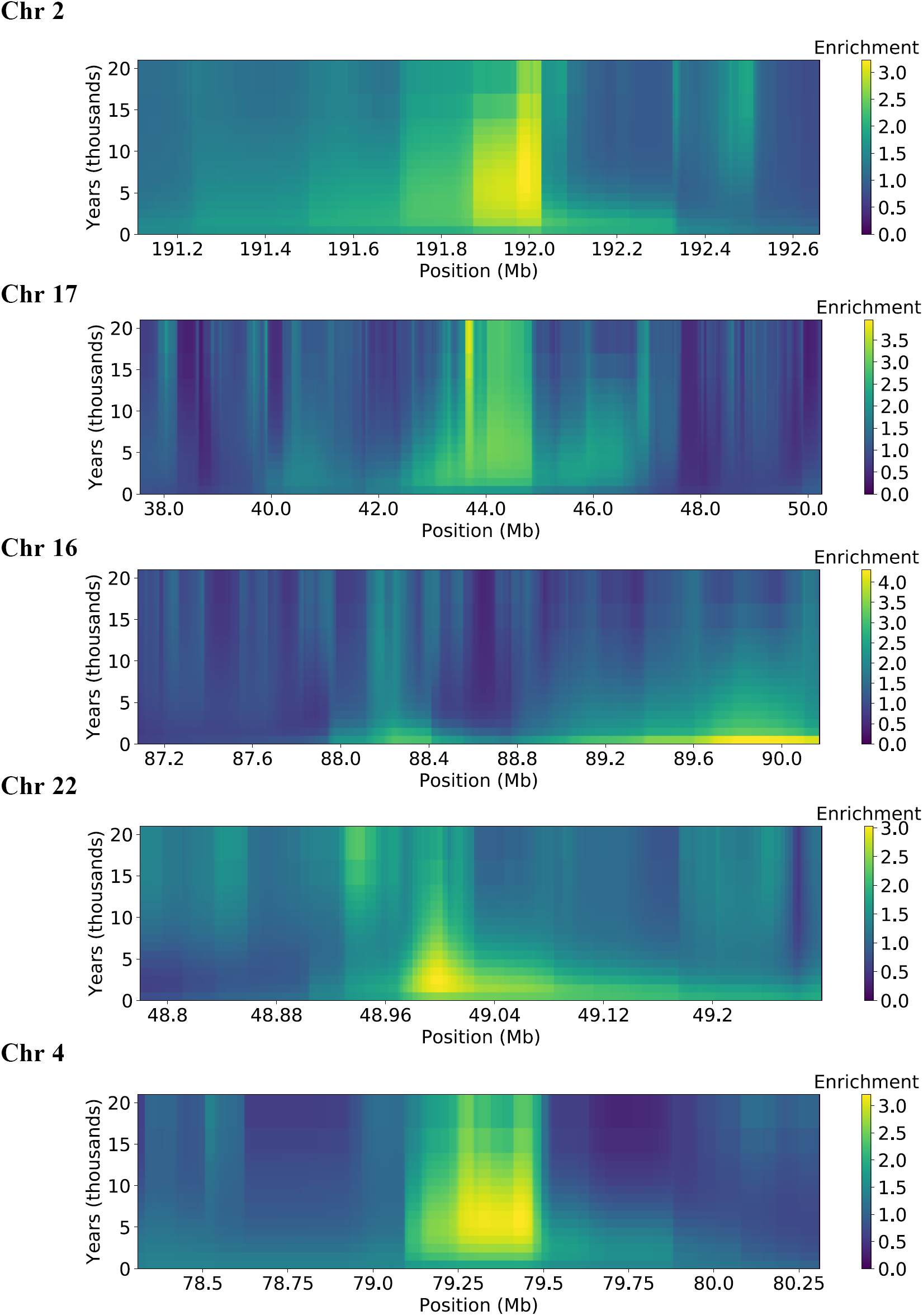
Enrichment of coalescence density in the past 20,000 years. At each site along the regions (horizontal axis) we plot the enrichment for the density of coalescence events in the past ∼20,000 years, computed as 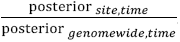. Time axes assumes a 30-year generation.

**Supplementary Figure 7.**
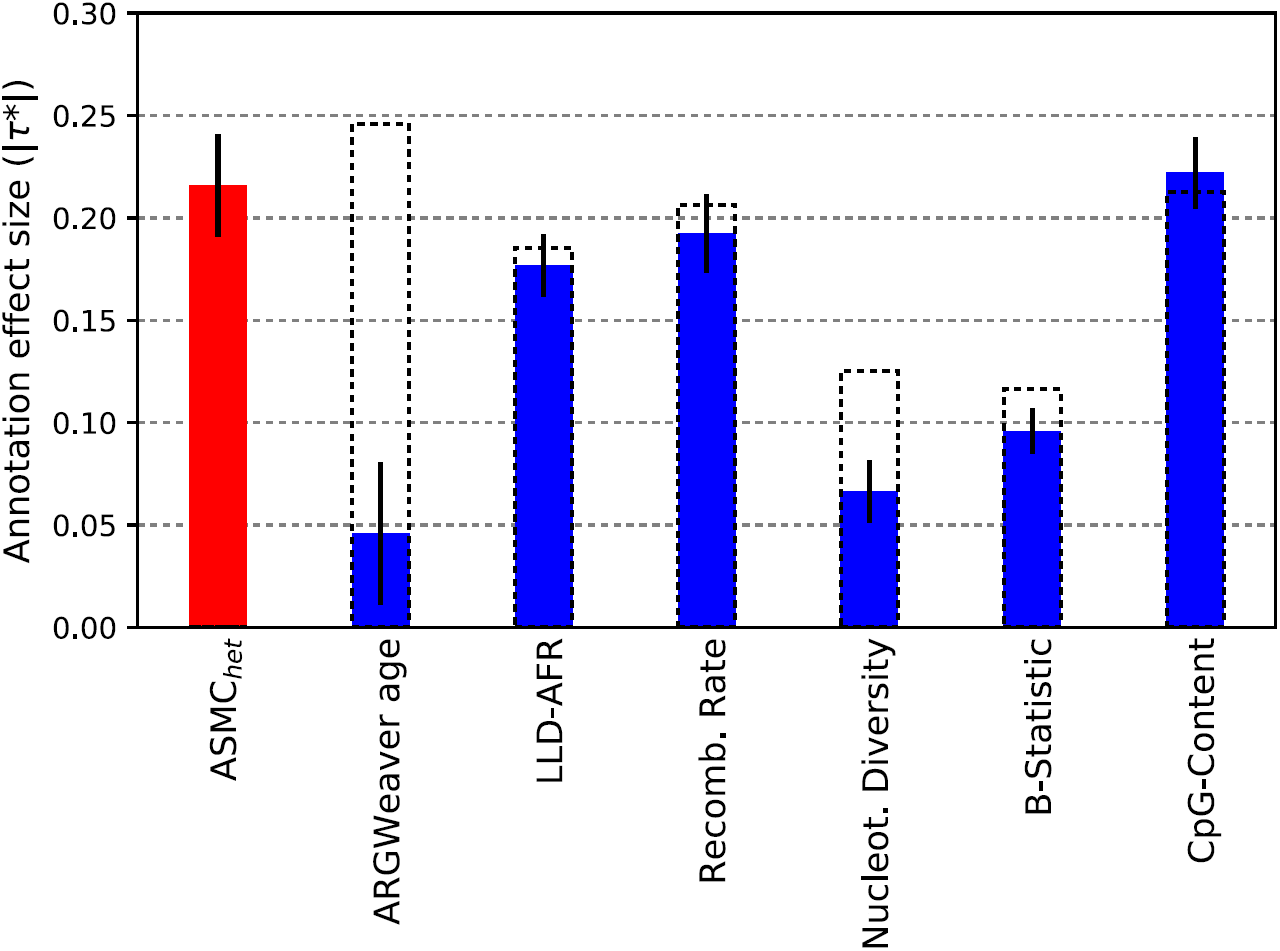
S-LDSC analysis of ASMC_het_ background selection annotation and disease heritability. We built an annotation, ASMC_het_, reflecting the average coalescence time for heterozygous individuals (i.e. chromosomes carrying discordant alleles) at each site. As for the ASMC_avg_ annotation, ASMC_het_ is quantile normalized using 10 MAF bins. ASMC_het_ is expected to be proportional to the age of polymorphic alleles in the sample. Consistent with this expectation, in a joint S-LDSC analysis using the ASMC_het_ annotation and the baselineLD model, we observed that the meta-analyzed *τ*\* for the quantile normalized ARGWeaver allele age annotation was reduced from 0.250 (s.e. 0.012) to −0.046 (s.e. 0.018).

**Supplementary Figure 8.**
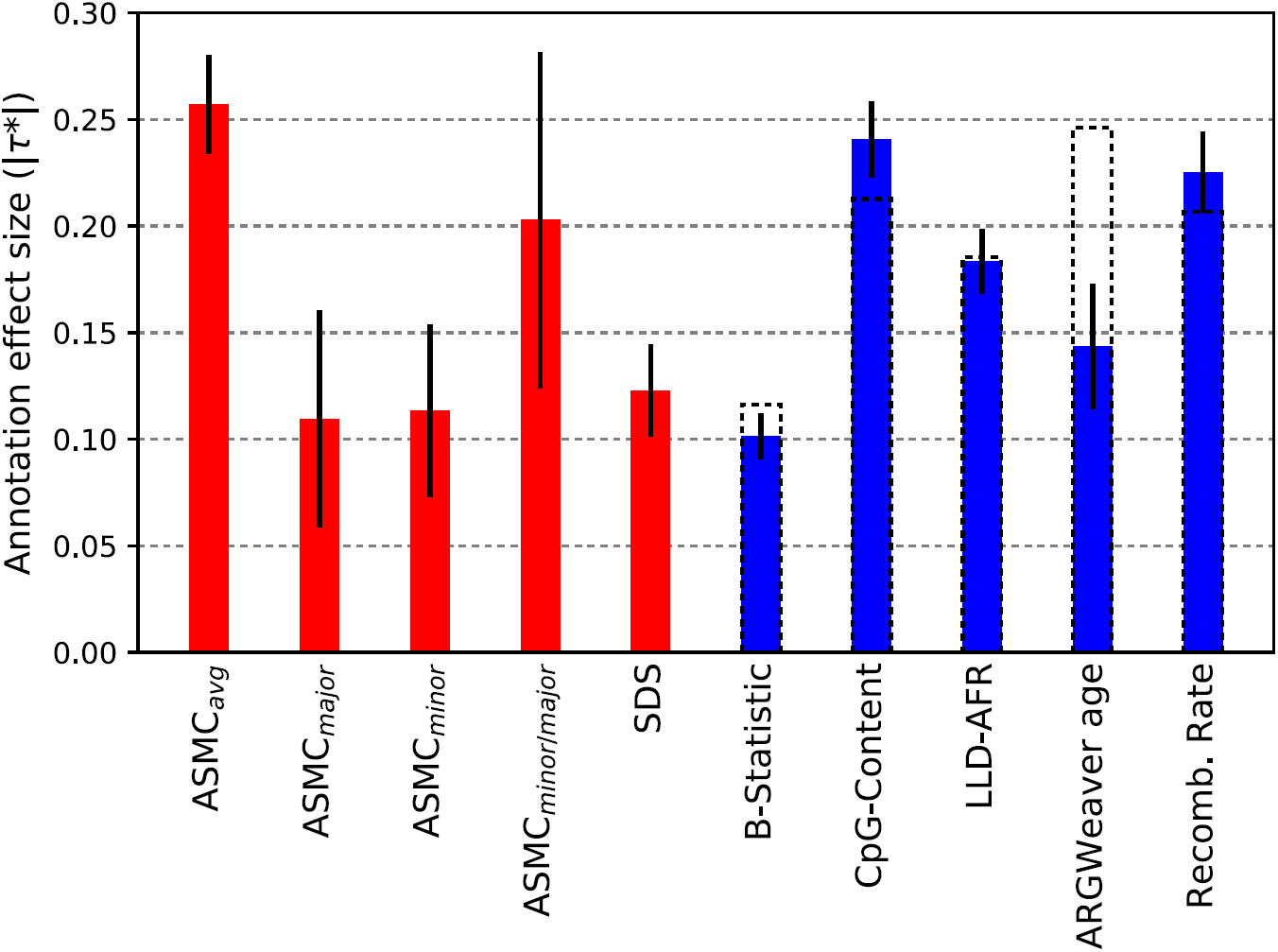
S-LDSC analysis of several annotations related to background selection. We built several annotations related to average coalescent time at each site, conditioning on the allele present on each analyzed chromosome from the GoNL data set. In addition to the ASMC_avg_ annotation (see **Online Methods**), we computed average coalescence time for carriers of a minor allele (ASMC_minor_), carriers of a major allele (ASMC_major_), and an annotation containing the value of log(T_minor_/T_major_) at each site, i.e. the logarithm of the ratio of average coalescence time for individuals carrying a minor allele and individuals carrying a major allele (ASMC_minor_/_major_). All annotations were quantile normalized with respect to 10 MAF bins, as done for the ASMC_avg_ annotation. We performed a joint S-LDSC analysis including these annotations, the SDS annotation from ref. ^36^, and all annotations from the baselineLD model, excluding the nucleotide diversity annotation, whose effects are subsumed by the ASMC_avg_ annotation (see Figure 4). ASMC_het_ was also excluded, as it was subsumed by ASMC_avg_. We report |*τ*\*| values meta-analyzed across 20 independent traits. Dashed lines for the baselineLD annotations represent meta-analyzed |*τ*\*| values in a joint S-LDSC analysis that does not include annotations represented in red.

**Supplementary Figure 9.**
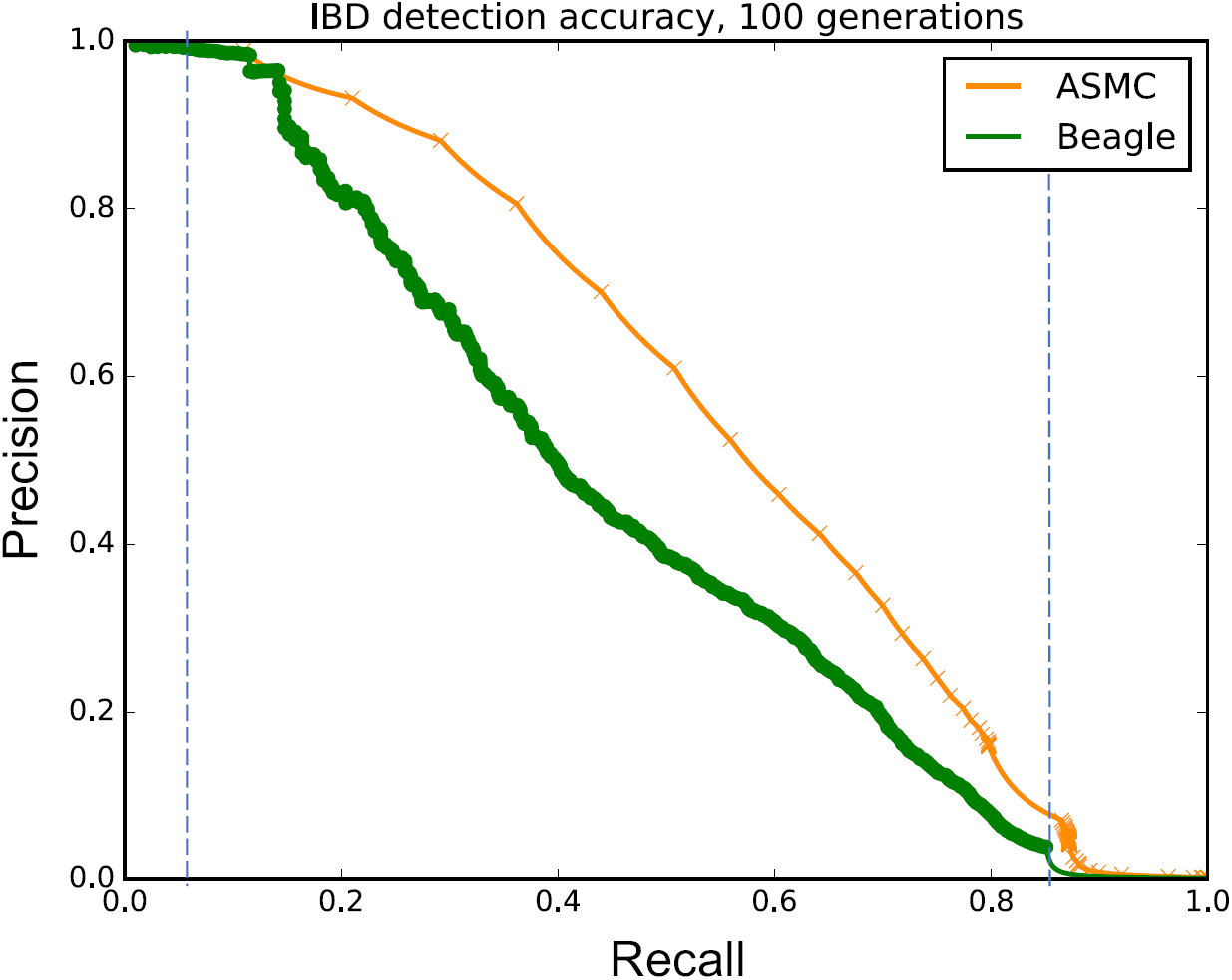
Illustration of auPRC measure for IBD detection accuracy. We measured accuracy of IBD detection for ASMC and Beagle using the area under the precision-recall curve (auPRC) for both programs. For both methods, recall can only be estimated within a limited precision range, due to the time-discretization used by ASMC, and the limited range of LOD-score thresholds allowed by Beagle. We thus compare the auPRC within the region where the precision and recall of both methods can be measured. In this example, ASMC’s recall can be measured for values greater than 0.05, while Beagle’s recall can be measured for values smaller than 0.85. We thus compare the auPRC for the two methods in the range [0.05, 0.85] (blue vertical lines). Interpolation between pairs of observed precision/recall values was obtained using the method of ref. ^80^.

**Supplementary Figure 10.**
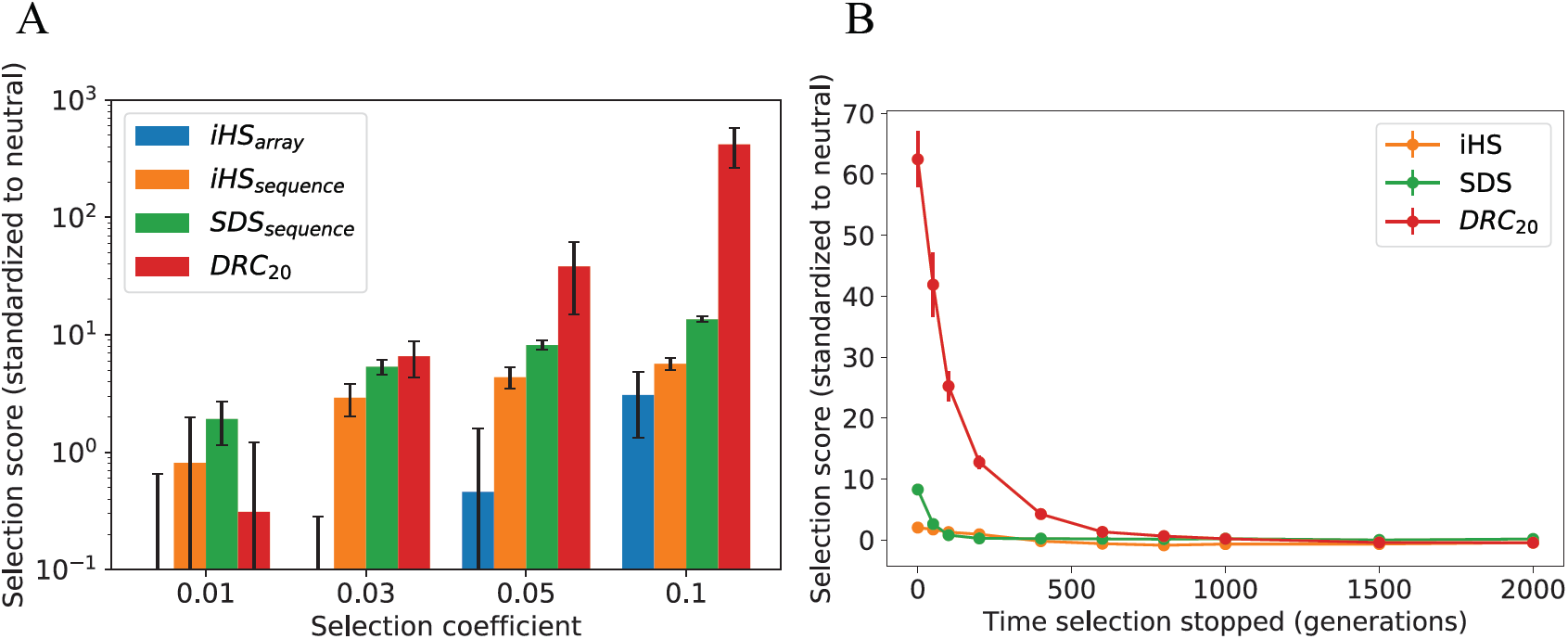
Selection simulations for DRC_20_. (A) Simulation of different strengths of recent positive selection starting 200 generations in the past: iHS score^35^ run on array data (iHSarray); iHS score^35^ on causal variant from sequencing data (iHS_sequence_); SDS score^36^ on causal variant from sequencing data (SDS_sequence_); DRC_20_ score on array data. Scores of each method are standardized with respect to corresponding scores obtained in neutral simulations. Bars indicate standard deviations. (B) Specificity to recent past for iHS and SDS run son sequencing data, and for DRC_20_. Simulation of selection starting at time -∞ stopping at the specified generation, followed by neutral drift. Error bars represent standard errors.

**Supplementary Figure 11.**
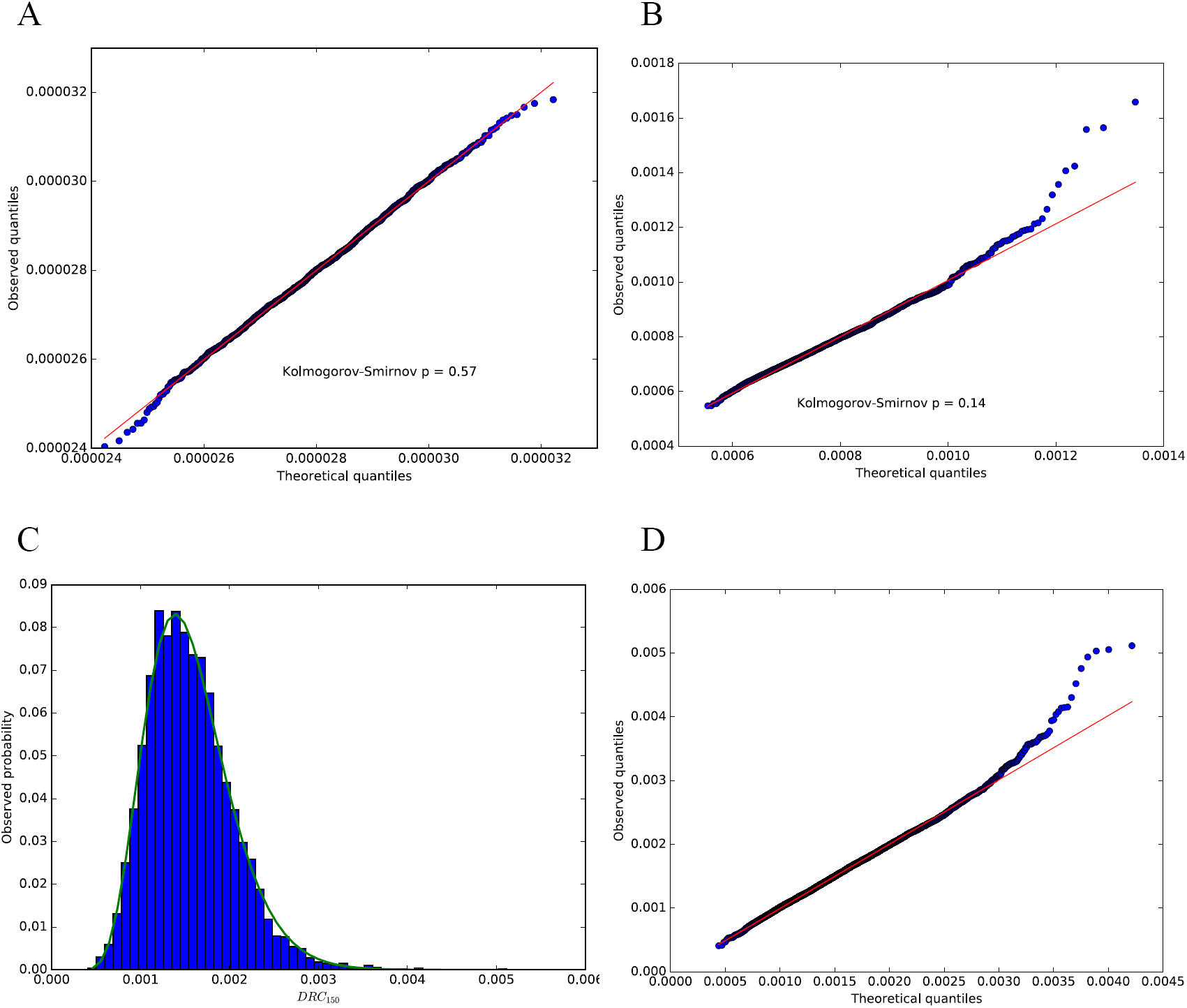
Empirical null model. (A) QQ plot for the DRC_20_ statistic in 2,000 independent neutral coalescent simulations using the European demographic model of Tennessen et al.^19^. (B) QQ plot for the DRC_150_ statistic in 2,000 independent neutral coalescent simulations using the European demographic model of Tennessen et al.^19^. (C) Empirical distribution and Gamma-fit for the DRC_150_ statistic in the putatively neutral portion of the genome in the UKBB data set (11,221 observations from 0.05 cM windows). (D) QQ plot for the DRC_150_ statistic in the putatively neutral portion of the genome in the UKBB data set (11,221 observations from 0.05 cM windows). All models are fit using a Gamma distribution with shape, location and scale parameters, using Python’s Scipy library (see URLs).

**Supplementary Figure 12.**
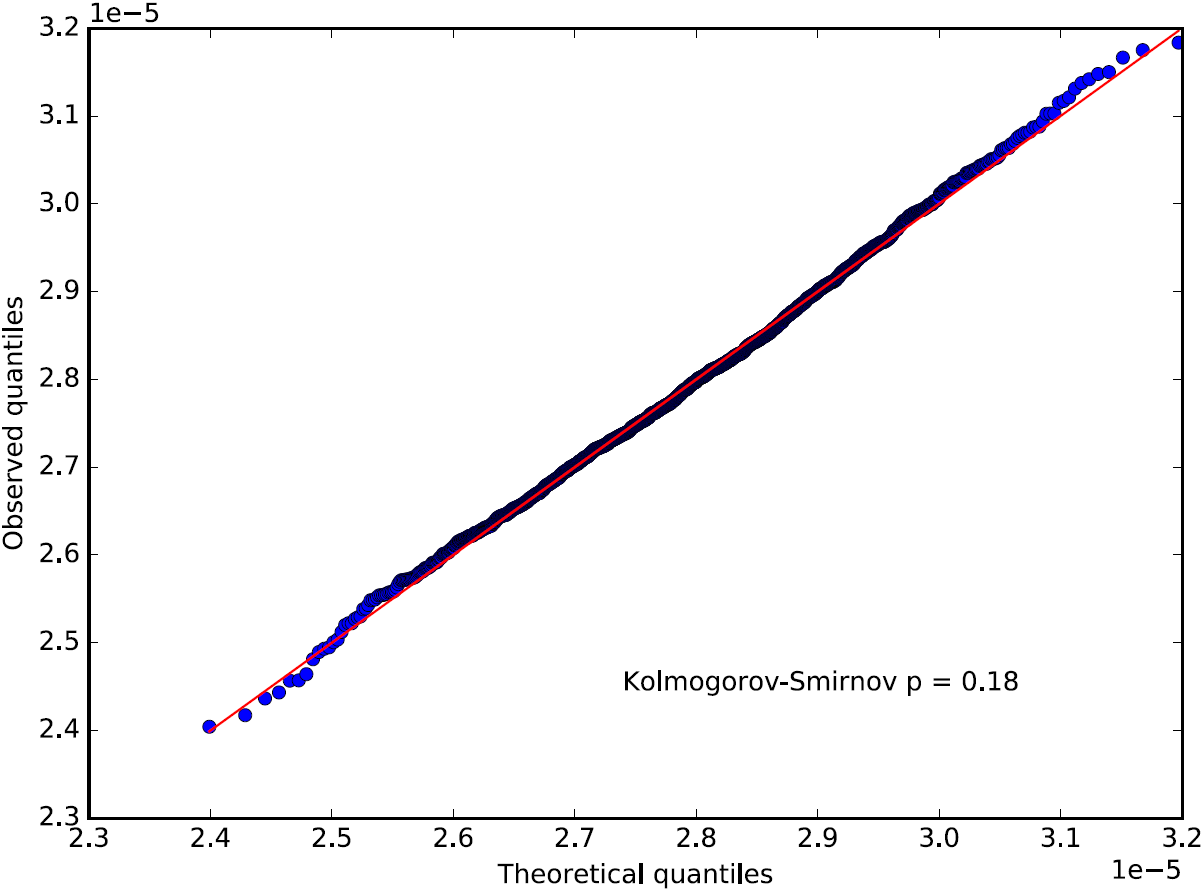
Empirical null model for the DRC_20_ statistic. QQ plot for the DRC_20_ statistic in 2,000 independent neutral coalescent simulations using the European demographic model of Tennessen et al.^19^, fit using a Normal distribution.

**Supplementary Table 1.** IBD detection. We report the difference in accuracy between ASMC-and Beagle-based IBD detection. IBD regions are defined using several time thresholds. We report the percent improvement for the area under the precision-recall curve (auPRC) of ASMC over Beagle. For both methods, precision can only be estimated within a limited recall range, due to the time-discretization used by ASMC, and the limited range of LOD-score thresholds allowed by Beagle. We thus compare the auPRC within the region where the precision and recall of both methods can be measured (“Average precision range” column, also see Supplementary Figure 9). Numbers in round brackets represent standard errors.

| IBD time threshold | Average recall range | auPRC within recall range | | Percent auPRC improvement:<br>$100 \times (\text{auPRC}_{ASMC} / \text{auPRC}_{Beagle} - 1)$ |
| --- | --- | --- | --- | --- |
|  |  | ASMC | Beagle |  |
| 25 | [0.26, 0.98] | 0.44 (0.01) | 0.38 (0.01) | 17.79 (2.18) |
| 50 | [0.12, 0.95] | 0.54 (0.01) | 0.44 (0.01) | 21.71 (1.22) |
| 75 | [0.07, 0.90] | 0.54 (0.01) | 0.44 (0.01) | 22.66 (1.11) |
| 100 | [0.04, 0.86] | 0.52 (0.00) | 0.43 (0.00) | 21.47 (0.95) |
| 150 | [0.02, 0.77] | 0.49 (0.00) | 0.41 (0.00) | 19.04 (0.81) |
| 200 | [0.01, 0.70] | 0.46 (0.00) | 0.39 (0.00) | 17.58 (0.53) |
| 400 | [0.00, 0.49] | 0.37 (0.00) | 0.31 (0.00) | 17.78 (0.39) |
| 600 | [0.00, 0.36] | 0.29 (0.00) | 0.24 (0.00) | 19.99 (0.37) |

**Supplementary Table 2.** Effects of demographic model misspecification. We simulated batches of 300 haploid samples from the first 30Mb of a human Chromosome 2 and a European demographic model, and ran ASMC using 160 discretization intervals (see **Online Methods**, Discretization Intervals). ASMC was ran assuming a constant effective population size of 10,000 diploid individuals, rather than the European model used to generate the data. We report percent difference in accuracy (RMSE and r^2^), compared to using the appropriate demographic model. We observed an increase in RMSE error compared to ASMC analysis using the correct demographic model, and no significant difference in r^2^. Numbers in round brackets indicate standard errors from 10 independent simulations.

|  | **% difference when using**<br>**wrong demographic model** |
| --- | --- |
| RMSE of posterior mean estimate of TMRCA | +29.18 (1.21) |
| RMSE of maximum-a-posteriori estimate of TMRCA | +82.29 (0.99) |
| $r^2$ of posterior mean estimate of TMRCA | -0.43 (0.49) |
| $r^2$ of maximum-a-posteriori estimate of TMRCA | +1.28 (1.29) |

**Supplementary Table 3.** IBD detection when ASMC demographic model is incorrect. We report the difference in accuracy between ASMC-and Beagle-based IBD detection. IBD regions are defined using several time thresholds. We report the percent improvement for the area under the precision-recall curve (auPRC) of ASMC over Beagle. For both methods, precision can only be estimated within a limited recall range, due to the time-discretization used by ASMC, and the limited range of LOD-score thresholds allowed by Beagle. We thus compare the auPRC within the region where the precision and recall of both methods can be measured (“Average precision range” column, also see **Supplementary Figure 9**). Numbers in parethesis represent standard errors. Data were simulated under a European demographic model, but ASMC was run assuming a constant effective population size of 10,000 diploid individuals. This had negligible effects on accuracy, although the TMRCA bias introduced by this model misspecification slightly shifted the average precision range where ASMC’s AuPRC could be measured.

| IBD time threshold | Average recall range | auPRC within recall range | | Percent AuPRC improvement:<br>$100 \times (\text{auPRC}_{ASMC} / \text{auPRC}_{Beagle} - 1)$ |
| --- | --- | --- | --- | --- |
|  |  | ASMC | Beagle |  |
| 25 | [0.32, 0.98] | 0.38 (0.01) | 0.32 (0.01) | 19.83 (2.92) |
| 50 | [0.15, 0.94] | 0.51 (0.01) | 0.41 (0.01) | 24.80 (1.40) |
| 75 | [0.08, 0.90] | 0.53 (0.01) | 0.42 (0.01) | 26.72 (1.55) |
| 100 | [0.05, 0.85] | 0.53 (0.01) | 0.42 (0.01) | 26.94 (1.09) |
| 150 | [0.03, 0.77] | 0.51 (0.00) | 0.40 (0.00) | 25.49 (0.80) |
| 200 | [0.02, 0.70] | 0.48 (0.00) | 0.39 (0.00) | 23.79 (0.44) |
| 400 | [0.01, 0.49] | 0.37 (0.00) | 0.31 (0.00) | 18.88 (0.49) |
| 600 | [0.00, 0.36] | 0.28 (0.00) | 0.24 (0.00) | 17.68 (0.58) |

**Supplementary Table 4.** Effects of noise in the recombination rate map. To mimic inaccuracies in the genetic map we simulated data using a human recombination map, and ran ASMC using a map with added noise. The recombination rate between each pair of contiguous markers in the map was altered by randomly adding or subtracting a specified percentage of its true value (% noise). We report accuracy using RMSE and r^2^. RMSE is measured between true and inferred TMRCA at each site, and “RMSE %” refers to the percent difference in RMSE between TMRCA inferred in SNP array data (UKBB density) using the indicated genetic map and TMRCA inferred in WGS data using the correct genetic map. Error attained using the true map is reported at the top for comparison. r^2^ indicates squared correlation between true and inferred average TMRCA in the simulated region. Each simulation involved a single batch of 300 haploid samples from the first 30Mb of a human Chromosome 2 and a European demographic model. ASMC was run using 160 discretization intervals (see Online Methods, Discretization Intervals).

| Map type | MAP RMSE % | Post. mean RMSE % | MAP $r^2$ | Post. Mean $r^2$ |
| --- | --- | --- | --- | --- |
| True map | +49.46 | +45.15 | 0.85 | 0.90 |
| 10% noise | +51.74 | +48.95 | 0.85 | 0.88 |
| 20% noise | +50.62 | +48.08 | 0.86 | 0.89 |
| 30% noise | +51.16 | +46.94 | 0.84 | 0.90 |
| 40% noise | +51.74 | +45.55 | 0.85 | 0.89 |
| 50% noise | +53.89 | +50.61 | 0.85 | 0.89 |
| 60% noise | +53.95 | +50.55 | 0.85 | 0.88 |
| 70% noise | +50.09 | +49.36 | 0.84 | 0.89 |
| 80% noise | +53.70 | +52.18 | 0.84 | 0.87 |
| 90% noise | +55.72 | +57.21 | 0.83 | 0.87 |
| 100% noise | +53.25 | +52.76 | 0.83 | 0.88 |

**Supplementary Table 5.** Effects of the number of time discretization intervals. We estimated coalescence times at each locus within 30Mb regions simulated using the standard setup using either the maximum-a-posteriori (MAP) or the posterior mean of the inferred coalescence distributions. In each simulation, we run ASMC using a different number of discretization intervals, which are chosen such that the coalescence distribution is expected to be uniform in all intervals (see Online Methods, Discretization intervals). For each value, we report the percent difference in RMSE accuracy between coalescence times inferred in SNP array data and WGS data, and the r^2^ between true and inferred average TMRCA in the regions.

| **Discretization intervals** | **MAP RMSE %** | **Post. mean RMSE %** | **MAP $r^2$** | **Post. mean $r^2$** |
| --- | --- | --- | --- | --- |
| 25 | +11.47 | +0.40 | 0.85 | 0.89 |
| 50 | +19.80 | -1.42 | 0.85 | 0.90 |
| 100 | +33.63 | +0.04 | 0.85 | 0.90 |
| 200 | +46.86 | +0.79 | 0.83 | 0.89 |
| 400 | +58.29 | +1.63 | 0.81 | 0.88 |

**Supplementary Table 6.**
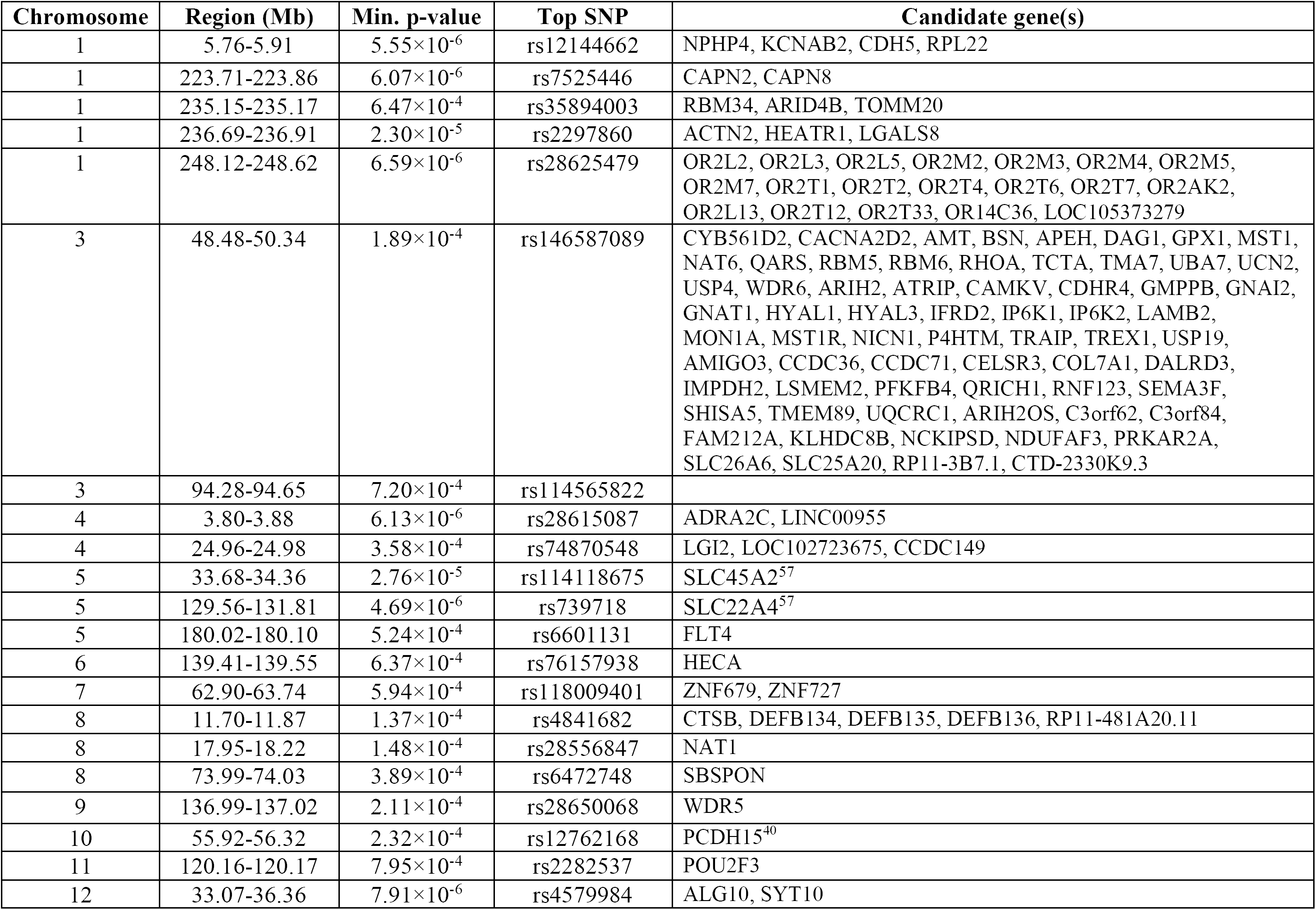
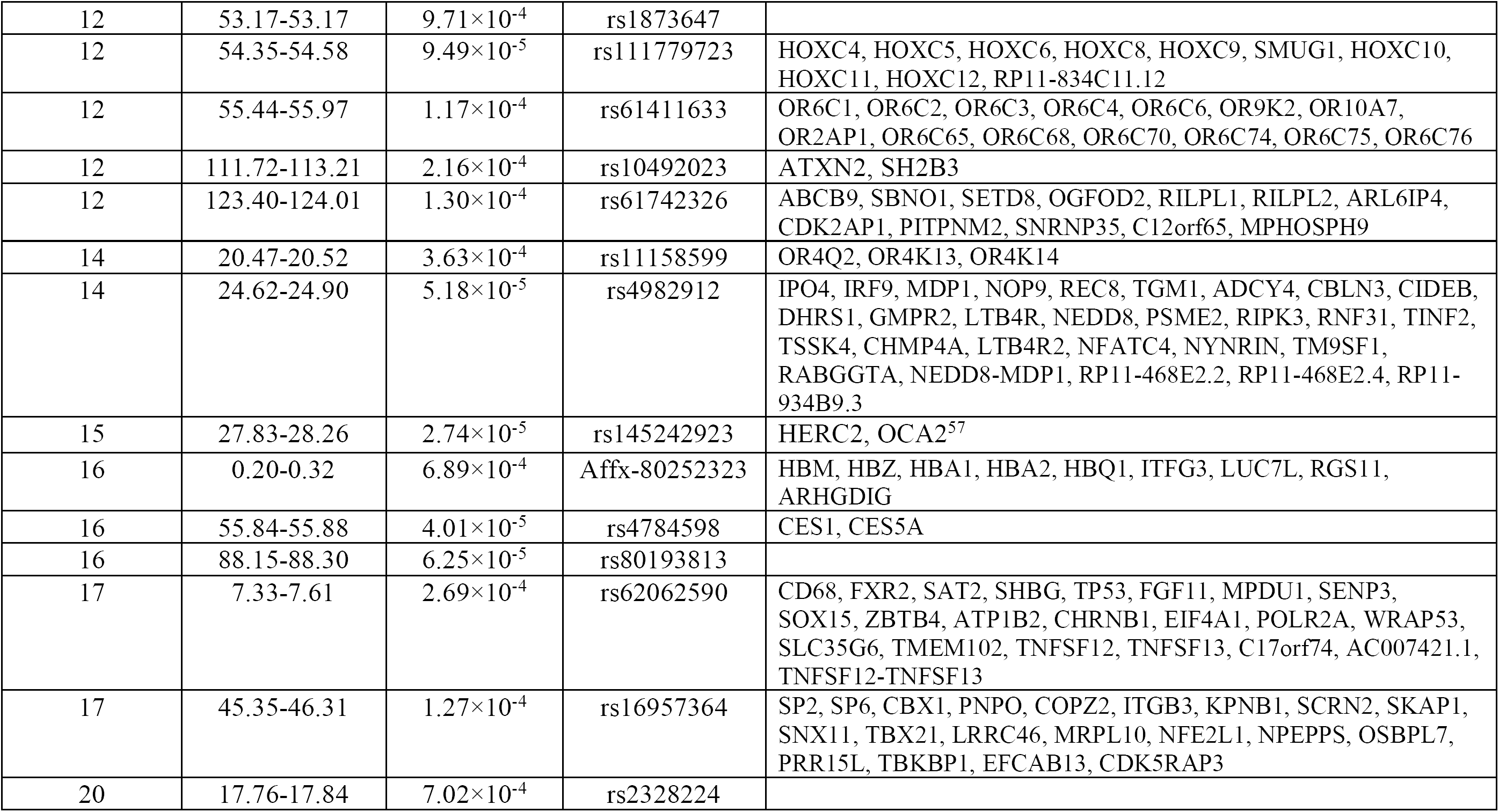
Suggestive selection loci. We report loci under suggestive selection (p < 10”^4^), as well as additional loci with elevated values of the DRC_150_ statistic in the UK Biobank data set (10^−4^ < p < 10^−3^).

**Supplementary Table 7.** Correlation between ASMC_avg_ and other annotations from the baselineLD model.

| **baselineLD annotation** | **Correlation with $ASMC_{avg}$ (r)** |
| --- | --- |
| B-statistic<sup>59</sup> | -0.28 |
| CpG content | 0.03 |
| Recombination rate | 0.07 |
| LLD-Africa<sup>61</sup> | 0.08 |
| ARGWeaver allele age<sup>6</sup> | 0.26 |
| Nucleotide diversity | 0.50 |

**Supplementary Table 8.** Traits analyzed in S-LDSC analysis. We report phenotype name (and reference), number of samples in the study, Z-score for the trait’s heritability, and URL (if summary statistics are publicly available).

| Phenotype | N | h<sup>2</sup> Z-score | URL |
| --- | --- | --- | --- |
| Age at menarche (UKBB) | 74,944 | 18.27 | . |
| Age at menopause (UKBB) | 44,410 | 9.87 | . |
| Anorexia<sup>87</sup> | 32,143 | 9.58 | http://www.med.unc.edu/pgc/downloads/ |
| Autism spectrum<sup>88</sup> | 10,263 | 8.72 | http://www.med.unc.edu/pgc/files/resultfiles/pgcasdeuro.gz |
| Blood pressure, diastolic (UKBB) | 134,011 | 24.12 | . |
| BMI<sup>89</sup> | 122,033 | 17.45 | http://www.broadinstitute.org/collaboration/giant/index.php/GIANT_consortium_data_files |
| Coronary artery disease<sup>90</sup> | 77,210 | 8.47 | http://www.cardiogramplusc4d.org/ |
| Crohn's disease<sup>91</sup> | 20,883 | 10.34 | http://www.ibdgenetics.org/downloads.html |
| Eczema (UKBB) | 145,416 | 11.42 | . |
| Height (UKBB) | 145,368 | 29.29 | . |
| LDL<sup>92</sup> | 93,354 | 9.49 | http://www.broadinstitute.org/mpg/pubs/lipids2010/ |
| Lung FEV1/FVC ratio (UKBB) | 123,935 | 22.04 | . |
| Lung forced expiratory volume (UKBB) | 123,935 | 27.99 | . |
| Neuroticism<sup>93</sup> | 170,911 | 9.54 | http://ssgac.org/documents/ |
| Putamen volume<sup>94</sup> | 12,924 | 7.08 | http://enigma.ini.usc.edu/wp-content/uploads/E2_EVIS |
| Rheumatoid arthritis<sup>95</sup> | 37,681 | 10.20 | http://plaza.umin.ac.jp/yokada/datasource/software.html |
| Schizophrenia<sup>96</sup> | 70,100 | 21.82 | http://www.med.unc.edu/pgc/downloads/ |
| Smoking status (UKBB) | 145,227 | 19.70 | . |
| Systemic lupus erythematosus<sup>97</sup> | 14,267 | 6.37 | https://www.immunobase.org/downloads/protected_data/GWAS_Data/ |
| Years of education<sup>98</sup> | 126,559 | 11.97 | http://www.ssgac.org/ |

**Supplementary Table 9.** Percent heritability explained by SNPs within annotation quintiles. We performed a joint analysis of ASMCTMRCA and other annotations in the baselineLD model using S-LDSC, and estimated the fraction of heritability explained by SNPs in each quintile of an annotation. The highest ratio between largest and smallest mean quintile effects was observed for the ASMC_avg_ and nucleotide diversity annotations. The effect (measured using *τ*\*, see Figure 4B) of the nucleotide diversity annotation, however, is subsumed by the ASMC_avg_ annotation.

| Annotation | % of heritability for SNPs in each quintile |  |  |  |  | largest/smallest |
| --- | --- | --- | --- | --- | --- | --- |
|  | 1<sup>st</sup> | 2<sup>nd</sup> | 3<sup>rd</sup> | 4<sup>th</sup> | 5<sup>th</sup> |  |
| $ASMC_{avg}$ | 33.11 (0.53) | 26.05 (0.21) | 16.00 (0.39) | 16.05 (0.22) | 8.73 (0.51) | 3.79 |
| Nucleotide diversity | 31.53 (0.36) | 24.37 (0.15) | 20.40 (0.08) | 15.44 (0.16) | 8.31 (0.38) | 3.79 |
| LLD-Africa<sup>61</sup> | 29.12 (0.38) | 23.89 (0.13) | 20.45 (0.07) | 16.58 (0.16) | 9.94 (0.33) | 2.93 |
| ARGWeaver allele age<sup>6</sup> | 29.25 (0.68) | 25.43 (0.23) | 14.42 (0.31) | 20.16 (0.27) | 10.69 (0.63) | 2.74 |
| CpG content | 11.94 (0.21) | 16.76 (0.16) | 20.11 (0.13) | 22.09 (0.12) | 28.49 (0.40) | 2.39 |
| B-statistic<sup>59</sup> | 13.17 (0.30) | 15.91 (0.14) | 19.57 (0.09) | 22.77 (0.12) | 28.35 (0.38) | 2.15 |
| Recombination rate | 19.08 (0.29) | 19.87 (0.20) | 20.71 (0.19) | 22.02 (0.15) | 18.27 (0.62) | 1.04 |

**Supplementary Table 10.**
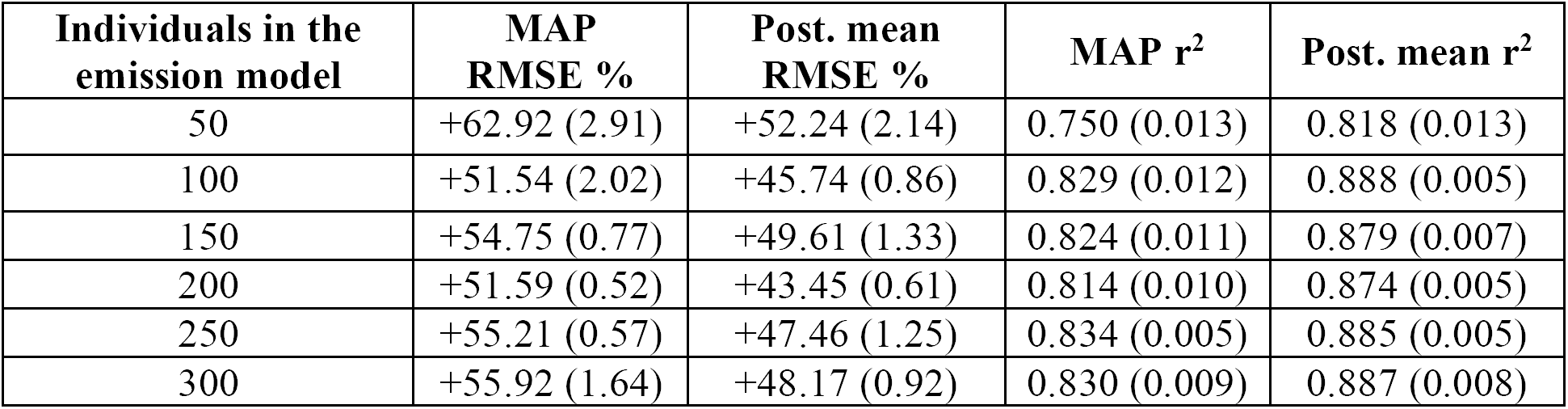
Effects of the number of samples used in the emission model. We simulated data using standard parameters, and measured accuracy of ASMC-inferred coalescence times using RMSE and r^2^ for either WGS and SNP array data. We estimated coalescence times at each locus using either the maximum-a-posteriori (MAP) or the posterior mean of the inferred coalescence distributions. In each simulation, we ran ASMC using 100 discretization intervals and a different number of samples to compute the CSFS in the emission model. For RMSE, we report the percent difference in accuracy between coalescence times inferred in SNP array data and WGS data. Better RMSE or r^2^ performance results for better use of allele frequency information via the CSFS emission model. We observed that the performance plateaus when using more than 100 samples in the CSFS. Approximate standard errors indicated in round brackets are computed using 5 independent simulations.

| **Individuals in the emission model** | **MAP RMSE %** | **Post. mean RMSE %** | **MAP $r^2$** | **Post. mean $r^2$** |
| --- | --- | --- | --- | --- |
| 50 | +62.92 (2.91) | +52.24 (2.14) | 0.750 (0.013) | 0.818 (0.013) |
| 100 | +51.54 (2.02) | +45.74 (0.86) | 0.829 (0.012) | 0.888 (0.005) |
| 150 | +54.75 (0.77) | +49.61 (1.33) | 0.824 (0.011) | 0.879 (0.007) |
| 200 | +51.59 (0.52) | +43.45 (0.61) | 0.814 (0.010) | 0.874 (0.005) |
| 250 | +55.21 (0.57) | +47.46 (1.25) | 0.834 (0.005) | 0.885 (0.005) |
| 300 | +55.92 (1.64) | +48.17 (0.92) | 0.830 (0.009) | 0.887 (0.008) |

**Supplementary Table 11.**
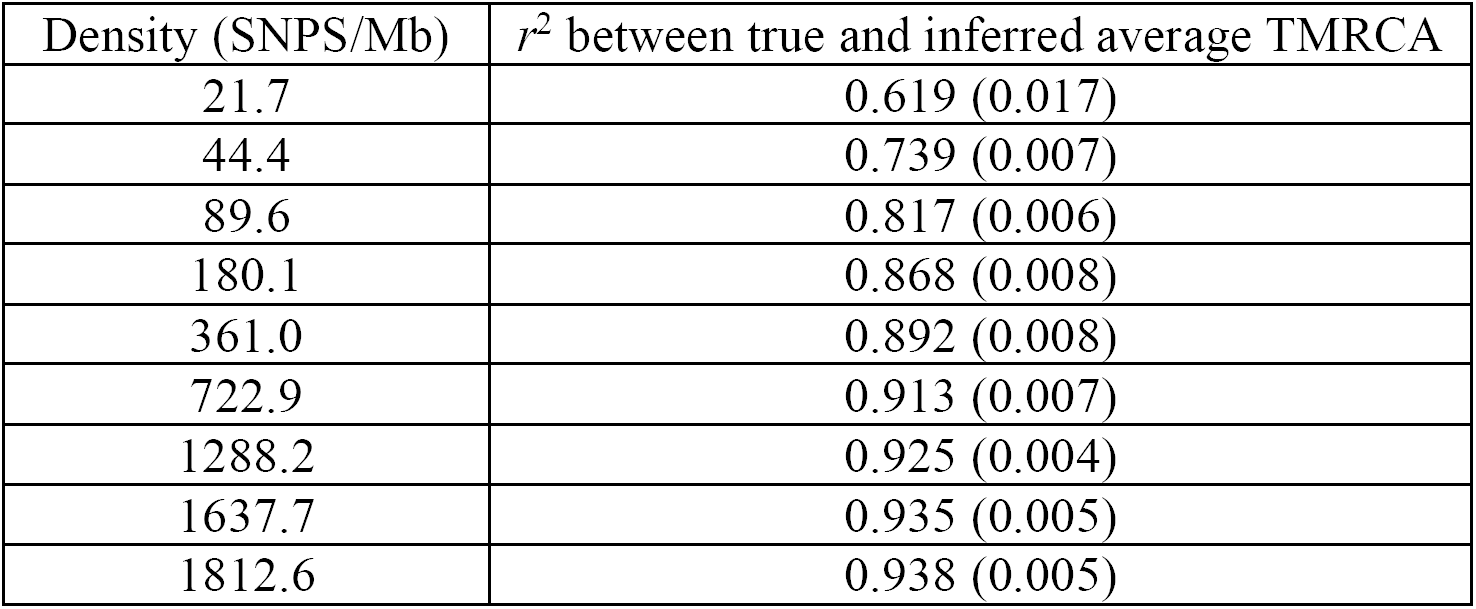
ASMC accuracy in coalescent simulations. Numerical values from Figure 1. Numbers in round brackets represent standard errors. The *r*^*2*^ attained by ASMC-seq using WGS data is 0.946 (0.017). Average SNP density observed in the UK Biobank data set was 225. TMRCA was inferred using the ASMC posterior mean coalescence time at each site within the simulated region.

| Density (SNPS/Mb) | $r^2$ between true and inferred average TMRCA |
| --- | --- |
| 21.7 | 0.619 (0.017) |
| 44.4 | 0.739 (0.007) |
| 89.6 | 0.817 (0.006) |
| 180.1 | 0.868 (0.008) |
| 361.0 | 0.892 (0.008) |
| 722.9 | 0.913 (0.007) |
| 1288.2 | 0.925 (0.004) |
| 1637.7 | 0.935 (0.005) |
| 1812.6 | 0.938 (0.005) |

**Supplementary Table 12.** Computational cost of ASMC. Numerical values from **Figure 2** and **Supplementary Figure 4**. Running times are extrapolated from those obtained in 5Mb long regions of WGS data, assuming a 3,235 Mb genome. Memory usage reflects analysis of a 5Mb region using WGS data from 100 haploid individuals. Numbers in round brackets represent standard errors.

| TMRCA intervals | Running time (seconds per genome) |  | Memory usage (Gb) |  |
| --- | --- | --- | --- | --- |
|  | ASMC-seq | SMC++ | ASMC-seq | SMC++ |
| 20 | 2.03 (0.05) | 211 (30) | 0.13 (0.0) | 0.37 (0.03) |
| 40 | 3.12 (0.06) | 1,150 (68) | 0.24 (0.0) | 0.83 (0.02) |
| 80 | 6.09 (0.21) | 6,310 (115) | 0.46 (0.01) | 1.63 (0.02) |
| 160 | 12.18 (0.41) | 43,879 (869) | 0.9 (0.01) | 3.67 (0.12) |
| 320 | 22.94 (0.97) | 347,385 (13,505) | 1.79 (0.03) | 9.88 (0.31) |
| 640 | 46.77 (0.77) | 2,649,817 (62,670) | 3.55 (0.06) | 21.67 (1.25) |

**Supplementary Table 13.** S-LDSC analysis of ASMC_avg_background selection annotation and disease heritability. Numerical values from **Figure 4**. (A) τ * value of the ASMC_avg_ annotation for 20 independent diseases and complex traits. (B) Absolute values of T* values (meta-analyzed across 20 independent diseases and complex traits) in joint analysis conditioned on baselineLD annotations. Values in round brackets represent standard errors. Numerical values for **Figure 4c** can be found in **Supplementary Table 9**

**A**
| Trait | $\tau^*$ |
| --- | --- |
| Rheumatoid Arthritis | -1.687 (0.233) |
| Crohns Disease | -1.452 (0.263) |
| LDL | -1.327 (0.229) |
| Eczema | -1.268 (0.189) |
| Age at Menopause | -1.111 (0.188) |
| Coronary Artery Disease | -1.093 (0.241) |
| Lupus | -1.082 (0.241) |
| Schizophrenia | -0.917 (0.078) |
| Diastolic | -0.865 (0.091) |
| Years of Education | -0.86 (0.131) |
| Height | -0.825 (0.094) |
| BMI | -0.759 (0.087) |
| FEV1FVC | -0.752 (0.089) |
| Smoking Status | -0.707 (0.088) |
| FVC | -0.686 (0.072) |
| Age at Menarche | -0.678 (0.107) |
| MeanPutamen | -0.629 (0.217) |
| Neuroticism | -0.536 (0.118) |
| Autism | -0.451 (0.192) |
| Anorexia | -0.25 (0.142) |
| Meta analysis | -0.807 (0.01) |

**B**
| Annotation | Annotation $\tau^*$ (meta-analysis) | Annotation $\tau^*$ (meta-analysis) not including $ASMC_{avg}$ in the model |
| --- | --- | --- |
| $ASMC_{avg}$ | -0.253 (0.010) | N/A |
| ARGWeaver allele age<sup>6</sup> | -0.133 (0.013) | -0.246 (0.012) |
| LLD-Africa<sup>61</sup> | -0.199 (0.008) | -0.185 (0.008) |
| Recombination rate | -0.223 (0.010) | -0.207 (0.010) |
| Nucleotide diversity | -0.001 (0.008) | -0.125 (0.008) |
| B-statistic<sup>59</sup> | 0.102 (0.006) | 0.116 (0.006) |
| CpG content | 0.220 (0.009) | 0.213 (0.009) |

## 1 Background

### 1.1 The pairwise sequentially Markovian coalescent (PSMC)

The pairwise sequentially Markovian coalescent (PSMC (Li & Durbin, 2011)) is a widely adopted coalescent-based hidden Markov model (HMM) that describes the ancestral relationship of a pair of haploid individuals at all sites along their genome. We provide a high-level description of this approach, upon which our model and several recent extensions have been built.

The vector of observations in the HMM is obtained from the genotypes of a pair of haploid individuals that are randomly sampled from a population. For a sequence of length ℓ, observations *x*_*i*_, *i* ∈ {1. ℓ}, have value 1 if the two individuals have discordant genotypes (they are heterozygous at the site) or 0 if they have identical genotypes (they are homozygous at the site). At each site along the genome, the hidden state *ti* ∈ {1*…d*} of the Markov chain represents the time to most recent common ancestor (TMRCA) of the pair of haploid individuals at site *i*. Time is measured in generations (or in coalescent units) and is discretized into a predefined set of d possible time intervals. The probability of observing a heterozygous site for the pair of individuals given their TMRCA is *t* is expressed as *P*(x = 1|*t*, μ) = 1 *e*^−2**μ*t*_*i*_^ where is the per generation, per base pair mutation rate, which is assumed to be constant along the genome and throughout time. Conversely, for a homozygous site, *P*(x = 0|*t, μ*) = 1 *e*^−2*μt*_*i*_^. The initial state probabilities for the HMM are obtained from the coalescent distribution induced by the effective size history of the population from which the two individuals were sampled. Transition probabilities between discrete TMRCA states along the genome are obtained using the sequentially Markovian coalescent (SMC) model, which provides a Markovian approximation to the coalescent process (McVean & Cardin, 2005) described as a sequence of recombination and coalescent events along the genome (Wiuf & Hein, 1999). Details of the transition model can be found in (Li & Durbin, 2011). The PSMC enables all usual applications of HMMs (Rabiner, 1989), including inferring the posterior probability of TMRCA at each site in the genome (posterior decoding), and learning the model’s hyperparameters, namely the population’s size history, mutation, and recombination rates.

### 1.2 Related work on coalescent HMMs

The CoalHMM model (Hobolth *et al.*, 2007) is one of the earliest coalescent HMMs, although its fundamental difference compared to the PSMC and derived approaches is that it operates at phylogenetic time scales, rather than population genetic time scales. The MSMC approach (Schiffels & Durbin, 2014), extended the PSMC to analysis of multiple haploid individuals. The hidden states of the MSMC model represent the time of the earliest coalescent event in the set of analyzed individuals, a modification that leads to increased insight into recent time scales. Another improvement of the MSMC over the PSMC is the use of the SMC’ model (Marjoram & Wall, 2006) in computing transition probabilities, which leads to increased accuracy compared to the SMC model (Hobolth & Jensen, 2014; Wilton *et al.*, 2015). When two individuals are analyzed, the MSMC approach reduces to the PSMC approach, though with the improved SMC’ transition model. The DiCal model (Sheehan *et al.*, 2013; Tataru *et al.*, 2014; Steinrucken *et al.*, 2015) is another coalescent HMM approach that enables simultaneous analysis of multiple samples, and explicit modeling of complex demographic scenarios. This approach relies on the conditional sampling distribution (CSD, (Paul *et al.*, 2011)), which approximates the full coalescent process by focusing on the conditional distribution of the n-th haploid individual given (*n —* 1) individuals have been observed. When two individuals are analyzed, the DiCal approach reduces to the PSMC model. The computational burden of both the MSMC and the DiCal approach limits their use to no more than ∼10 haploid individuals. The recently developed SMC++ method (Terhorst *et al.*, 2016), extends the PSMC approach by incorporating knowledge of the frequency of the analyzed genetic polymorphisms in the emission model the of HMM, effectively utilizing genotype information from multiple samples while computing posterior coalescent probabilities for a single pair of haploid individuals. To achieve this, the SMC++ approach crucially relies on the notion of “conditioned sample frequency spectrum” (CSFS, see section 2.1 for an overview, and (Terhorst *et al.*, 2016) for details). As in the MSMC approach, the transion model of the SMC++ provides an improvement over the PSMC’s approximation of the full coalescent process. The SMC++ adopts the conditional Simonsen-Churchil model (CSC) proposed in (Hobolth & Jensen, 2014), which is superior to the SMC’ approach, as it considers the possibility of multiple recombination events occurring between two sites without affecting the TMRCA for a pair of analyzed individuals.

### 1.3 Computational cost and phasing requirements of other methods

Standard computation of posterior probabilities via the forward-backward algorithm, which we will simply refer to as “posterior decoding” in the remainder of this note, has cost *𝒪* (*d*^*2*^*ℓ*) for d hidden states and an observation sequence of length ℓ (Rabiner, 1989). The standard forward-backward calculations adopted in the PSMC and MSMC methods therefore lead to *𝒪*(*d*^2^ℓ)computational cost to estimate posterior coalescent probabilities for a set of d discretized TMRCA intervals and a sequence of length *‘* base pairs. PSMC reduces computational costs by pooling sites in blocks of 100 base pairs, while MSMC uses precomputation and caching to improve run time. The DiCal method (Steinr ücken et al., 2015) uses a “locus-skipping” approach (Paul & Song, 2012), which enables running the forward-backward algorithm in time *O*(*d*^*2*^*ℓ*_*p*_), where *ℓ*_*p*_ is the set of loci that are polymorphic in the analyzed samples. This leads to substantial speed-ups, since usually ℓ ≫ ℓ_p_. A previous version of DiCal utilizes properties of the SMC model to reduce the run-time complexity of the forward-backward algorithm to *O*(*dℓ*) (Harris *et al.*, 2014). These approaches, however, are limited to use within the CSD model, which reduces to the SMC model when two haplotypes are analyzed. Compared to the SMC’ and the CSC model, the SMC provides a less accurate Markovian approximation of the coalescent (Hobolth & Jensen, 2014; Wilton *et al.*, 2015). The SMC++ approach, which utilizes the more accurate CSC model, implements a novel “locus-skipping” approach that enables computing the forward-backward dynamics in time *𝒪* (*d*^*3*^*ℓ*_*p*_).

The coalescent HMM approaches discussed thus far require the availabulity of accurate phasing information in order to perform TMRCA posterior decoding for haplotypes from distinct diploid individuals. Accurate computational phasing, however, cannot be achieved in modern sequencing data sets, particularly for rare variants. This often limits the application of coalescent HMM approaches to the maternal and paternal haplotypes within unphased diploid individuals, or results in noisy estimates of TMRCA distributions in the presence of pervasive phasing errors (Terhorst *et al.*, 2016). Although the SMC++ approach provides an effective way of pooling information from the genotype of multiple unphased individuals from a sample, TMRCA decoding for pairs of haplotypes sampled across different diploid individuals still requires access to phasing information.

## 2 The ascertained sequentially Markovian coalescent

Here, we develop the Ascertained Sequentially Markovian Coalescent (ASMC). The ASMC is most closely related to the SMC++ (Terhorst *et al.*, 2016), and makes the following methodological innovations:

- The ability to perform posterior decoding using a non-random subset of genomic variants, such as the subset of common variants that are genotyped using SNP array technologies.
- A new formulation of the forward-backward algorithm that requires *𝒪* (*dℓ*_*p*_) computation under the conditional Simonsen-Churchil transition model (Hobolth & Jensen, 2014).

These two advances enable performing high-troughput coalescent-based analysis of relatedness in large SNP array data sets, which are now widely available and often comprise several tens or hundreds of thousand samples. Furthermore, owing to recent advances in computational phasing algorithms (Loh *et al.*, 2016a; Loh *et al.*, 2016b; O’Connell *et al.*, 2016), large cohorts such as the UK Biobank can now be computationally phased with very high accuracy, with switch error rates in the order of 0.3% (one every ∼10 cM). This creates the possibility of analyzing coalescent times for potentially all pairs of haploid individuals in the sample, with negligible effects of phasing errors. The dramatic speedup achieved by ASMC also makes analysis of all pairs of available haploid genomes feasible in sequencing data sets, whenever high-quality phasing information is available.

### 2.1 ASMC emission

The emission model of a coalescent HMM approach for the inference of TMRCA in non-randomly ascertained genotype data, such as SNP array data, needs to tackle two key technical challenges, namely

1. The information content of the observed genotype data with respect to the coalescent time of the analyzed individuals is greatly reduced, as the vast majority of genotype variants are unobserved.
2. The set of observed variants are not randonly ascertained from the underlying sequencing variants. This ascertainment leads to significant bias in TMRCA inference if not accounted for.

To address these challenges, the ASMC adopts and extends the “conditioned sample frequency spectrum” (CSFS) model (Terhorst *et al.*, 2016). In addition to modeling allele sharing at each genomic site along the genome of the analyzed pair of individuals, as done in the PSMC approach, the CSFS enables taking into account the total number of individuals carrying each derived allele in a population sample. Modeling of allele frequencies using the CSFS allows to (1) increase the informativeness of the observations, enabling inference of TMRCA despite a substantial reduction in genotyped variants (2) remove biases due to frequency-based ascertainment, by explicitly modeling the probability of observing a variant in the data provided it is polymorphic at a given frequency in the analyzed sample.

The CSFS model can be briefly described as follows. Having obtained a set of (n + 2) haploid samples from a panmictic population with known demographic history, we denote 2 of these samples as “distinguished”, and the remaining *n* as “undistinguished”. Given that the pair of distinguished lineages coalesce at time τ at a site along the genome, the CSFS expresses the probability that exactly *d* out of the two distinguished individuals and *u* out of the n undistinguished individuals carry a mutated allele. We denote this probability as *CSFS(τ)*_*d,u*_, so that a *CSFS(τ)* is a 2 × n table where entry {d, u} corresponds to the probabily that d derived alleles are observed in the two distinguished samples, and *u* derived alleles are found in the *n* undistinguished samples (the value of n is dropped to simplify the notation). Details on the derivation of the CSFS for a given demographic model can be found in (Terhorst *et al.*, 2016). We note that in this paper we are mainly concerned with the task of decoding TMRCA along the genome of a pair of haploid individuals, and we will and adopt a demographic model inferred from previous analysis of whole-genome sequencing data.

Assume now that variants in the observed data set have been genotyped based on their frequency in a population sample, in other words, that the probability of observing a variant in the data can be expressed as *P*(*obs*|*d* + *u*). The *ascertained* conditioned site frequency spectrum is then obtained as *ACSFS(τ)*_*d*_,_*u*_ = *CSFS(r*)_*d,u*_ × *P*(*obs*|*d* + *u*) × *norm*, where *norm* is a normalizing constant such that 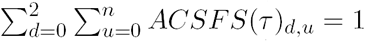. In practice, we need to estimate P*(obs*|d + u), and we do so by computing 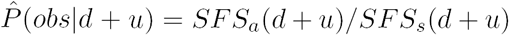, where *SFS*_*a*_*(x)* and *SFS*_*a*_*(x)* represent counts for the number of sites polymorphic in *x* individuals for a sample of size *n* + 2. Note that the normalization of *SFS*_*a*_(.) and *SFS*_*s*_(.), which should take into account terms related to e.g. the population mutation rate, is irrelevant, as these scaling constants vanish when the ACSFS is renormalized. To estimate the ascertained SFS_a_(-), we compute the sample frequency spectrum in the analyzed data. The sequence-level site frequency spectrum, *SFS*_*s*_(.) is obtained using the population demographic model, which is known and provided in input. The unconditioned site frequency spectrum may be obtained from the CSFS as 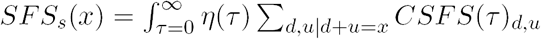 where *η* (*τ*) is the coalescent probability for the known demographic model at time τ.

### 2.2 ASMC transition

The transition model of a coalescent HMM dealing with sparsely ascertained data needs to account for the increased distance between observed markers. Observed variants in common SNP array data sets, for instance, are separated by several kilobases on average. The SMC transition model (McVean & Cardin, 2005) originally adopted in the PSMC approach (Li & Durbin, 2011) becomes particularly inaccurate in this setting, as it postulates that at most one recombination event may occur between two contiguous sites. Furthermore, the SMC assumes that any recombination event leads to a change in the value of the TMRCA, whereas the full coalescent model admits the possibility that a recombination event between two loci is followed by a coalescent event to the same lineage such that the TMRCA remains unchanged. This modeling limitation is mitigated in the improved SMC’ model (Marjoram & Wall, 2006), which allows for multiple recombination and coalescent events between two loci, and is adopted (though allowing for at most one recombination event) in the MSMC approach (Schiffels & Durbin, 2014). The ASMC transition model adopts the “conditional Simonsen-Churchil” model (CSC) described in (Hobolth & Jensen, 2014), also implemented in the SMC++ approach (Terhorst *et al.*, 2016). The CSC further improves modeling of recurring recombination and coalescent events between a pair of sites that are separated by large genetic distances, such as markers in SNP array data.

### 2.3 A general linear time forward-backward algorithm

Although several computational improvements have been proposed in previous coalescent HMM methods (see Section 1.3), further speed-ups are required for the analysis of all pairs of haploid samples in large data sets under the CSC model. We thus devise a new algorithm that enables performing forward-backward posterior calculations using the CSC transition model in time *𝒪 (dℓ*_*p*_), where *ℓ*_*p*_ is a set of observed loci for which we want to estimate TMRCA, and d is the number of discrete hidden TMRCA states. We start by introducing the Conditional Simonsen-Churchill model (Hobolth & Jensen, 2014), making use of the notation reported in Table 1.

**Table 1:**
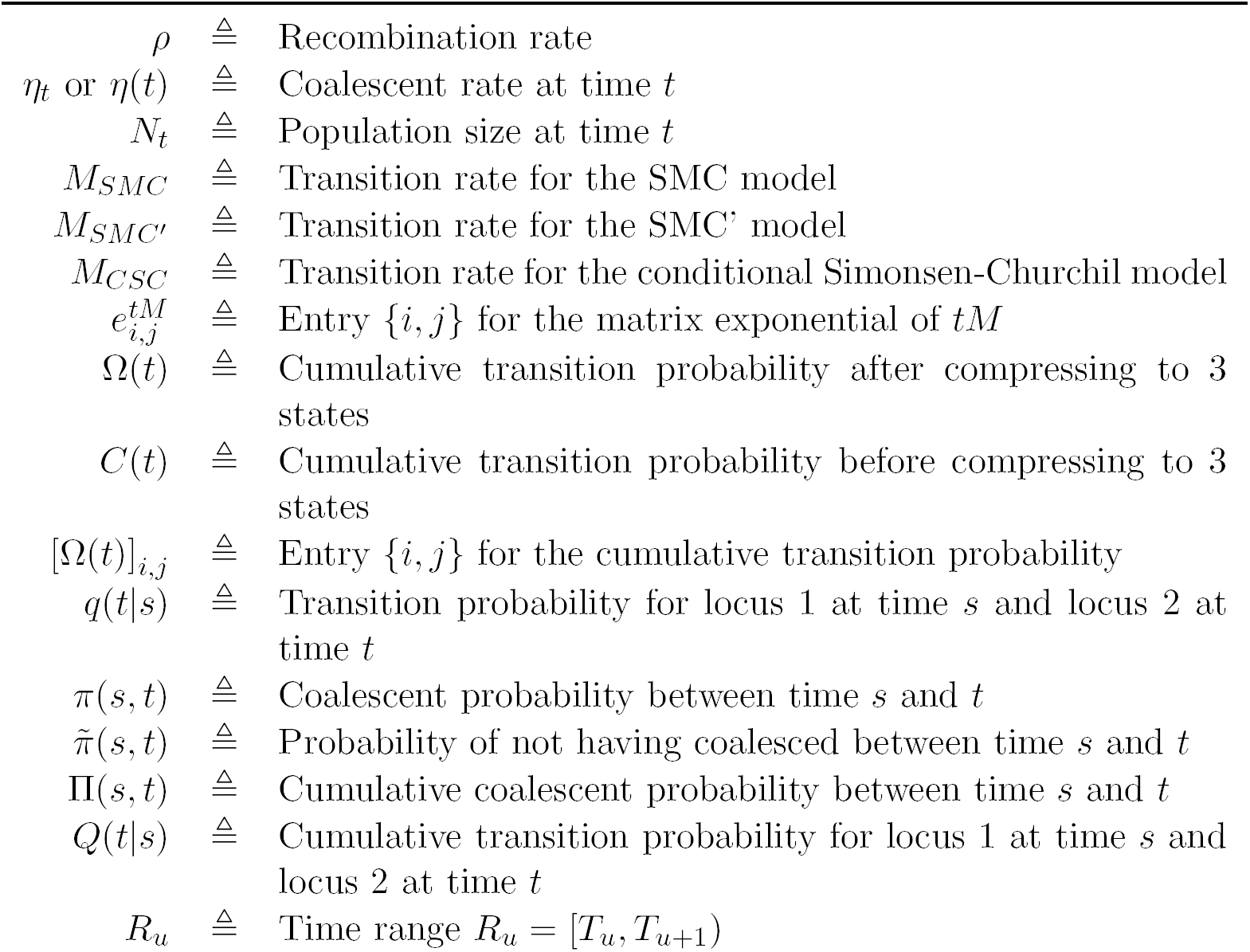
Table of notation for current section

#### 2.3.1 The conditional Simonsen-Churchill model

Consider two loci at recombination distance *ρ /2* in a population of constant size N, corresponding to a per-generation coalescent rate of *η* In (Hobolth & Jensen, 2014), the Markov chain of Figure 1 was used to descibe the distribution of ancestry at one locus conditional on the ancestry at the other locus. The transition matrix for this model is

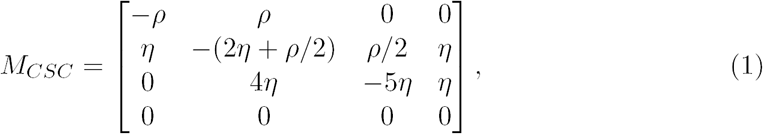

where each row and colum of the matrix represents one of the four states for which *t < s* (circles labeled with letters to the left of the vertical bar in Figure 1). Although the CSC model has four states, we will be mostly concerned with the probability that the Markov chain is in one of the three numbered states in Figure 1 that is, it will be irrelevant for our calculations whether at a given point in time the exact state of the chain is either state *B* or *C* within the dashed box. We thus define the matrix *Ω (t)*, whose first row is [*Ω* (*t*)]_1 •_ = [*C(t)AA,C(t)AB + C(t)*_*Ac*_,*C(t)AD*], where 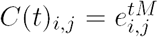 is the cumulative probability of transitioning from state *i* to state *j* after time *t*, for *i,j* ∈ *{A,B,C,D*}.

**Figure 1:**
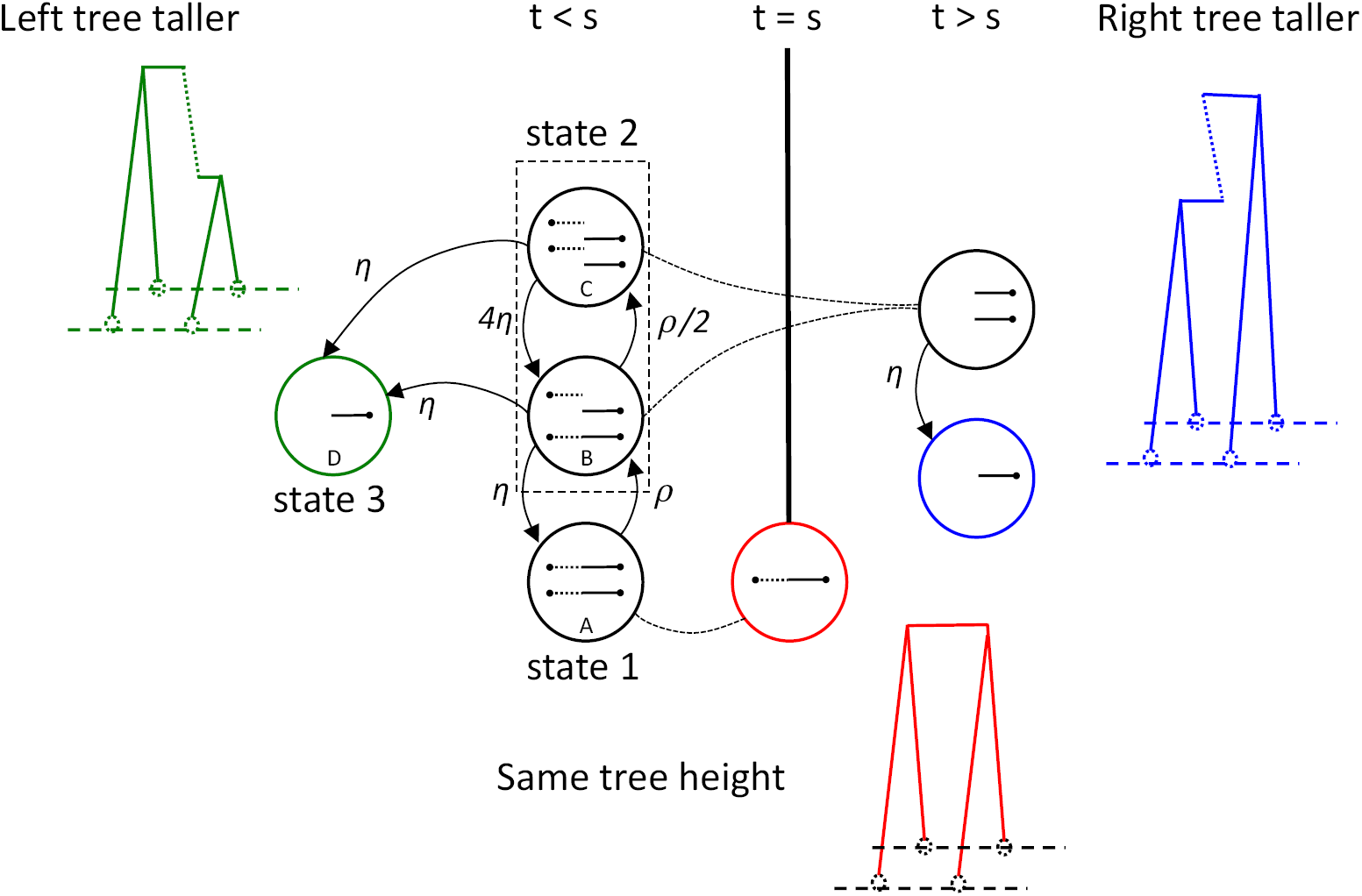
The conditional Simonsen-Churchil model (modified from Figure 1b of (Hobolth & Jensen, 2014)). Four relevant states from the full CSC model are labeled using letters within each circle.

For completion, we note that although we are mostly concerned with the CSC model, the discussion below also applies to the SMC and SMC’ models, which may be seen as special cases of the CSC where states *B* and *C* have been collapsed, with updated rate matrices

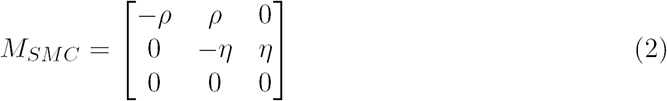

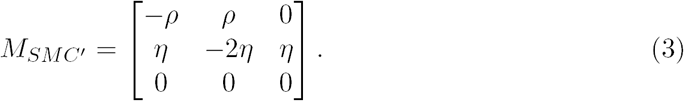

Note that *M*_*SMC*′_actually represents a process that is similar, but slightly different from the SMC’, as discussed in (Wilton *et al.*, 2015). Thus, [*Ω*(*t*)]_11_ will hold the probability that no recombination occurred from time 0 to time *t* or, for the SMC’ and CSC models, that at least one recombination occurred, but the lineages colasced back to state 1. [*Ω*(*t*)]_12_ respresents the probability that recombination occurred after time 0, but the lineages have not recoalesced back to state 1 or to a state such that the right tree has coalesced (state 3). [*Ω*(*t*)]_13_ represents the probability that the right tree is lower than the left tree, i.e. the two lineages coalesced at time *t < s.* Using these quantities, we can write the transition distribution for the height of the right tree, *t*, conditional on knowing the height of the left tree, *s* as

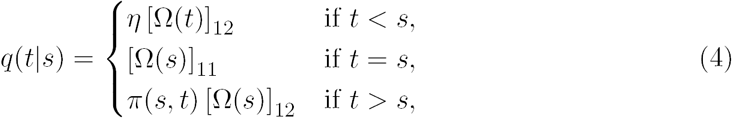

where *π* (*s,t*) is the coalescent probability between time *s* and *t*. This probability is computed as 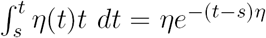 for a constant population size with coalescent rate *η*. Equation 4 is normalized, since

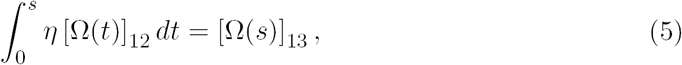

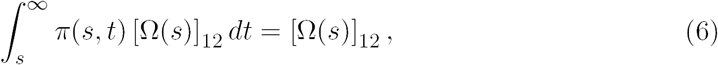

and[Ω(*s*)]_11_+[Ω(*s*)]_12_+[Ω(*s*)]_13_=1

##### 2.3.1.1 Piecewise constant demographic model

If the population size is piecewise constant, for each time period *k* ranging in *R*_*k*_ ∈ *[T*_*k*_,*Tk*_*+ i*_), there is a different transition rate matrix *Mk*. If *t* is contained in the interval *R*_*k*_, then the state matrix at time *t* can be computed as

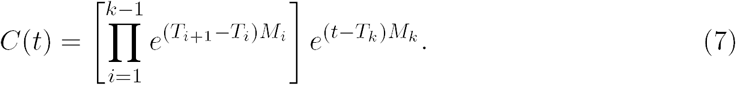

For a piece-wise constant model, the coalescent probability after time *s* can be similarly computed as

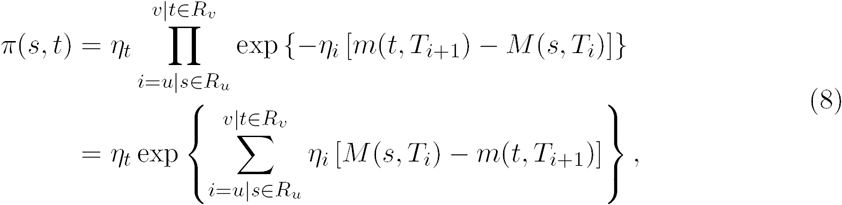

where *M*(…) and *m*(…) indicate maximum and minimum, respectively. The rate 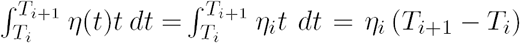 in the argument of the exponential should be substituted with the appropriate rate for inhomogeneous (e.g. exponential) models. We indicate the probability of not having coalesced at time *t* with

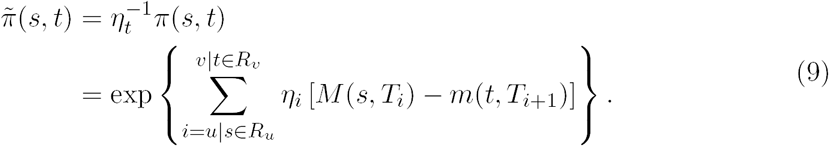

and the cumulative coalescent probability with

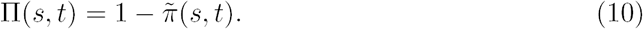

Using the quantities above, the transition probability for tree heights is still given by

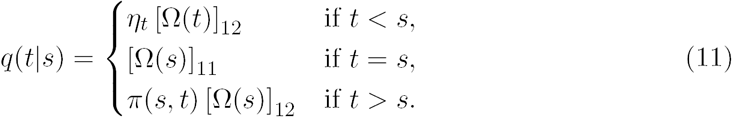

##### 2.3.1.2 Discretization

Using Equation 5, the cumulative transition probability is

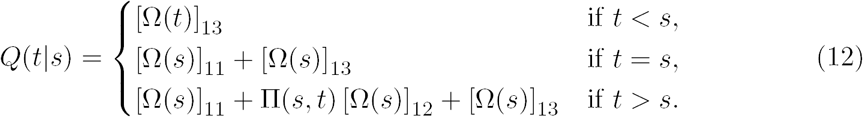

The probability of transitioning between time s and the time range *R*_*u*_ is then obtained as *Q*(*T*_*u*+1_|*s*) — *Q*(*T*_*u*_|*s*). The same approach can be used to further partition time in discrete states that do not necessarily correspond to population size changes. If we assume time has been discretized into *d* intervals, then we can obtain a transition matrix T such that entry *T*_*i,j*_ corresponds to the probability of transitioning from time interval i to time interval *j*. Each entry of the transition matrix is then obtained as *T*_*ij*_ = *Q(T*_*j+ 1*_*\s*_*i*_*) — Q(T*_*j*_+|s_*i*_), where we indicated the expected coalescent time within interval *R*_*i*_ as *s*_*i*_.

#### 2.3.2 Linear time computation of posterior coalescent times

We now describe a forward-backward algorithm to compute posterior coalescent probabilities in time *𝒪 (dℓ*_*p*_), where d is the number of discrete coalescent time intervals, and *ℓ*_*p*_ is the number of sites for which we wish to obtain TMRCA estimates (e.g. the set of observed sites). We use the notation reported in Table 2

**Table 2:**
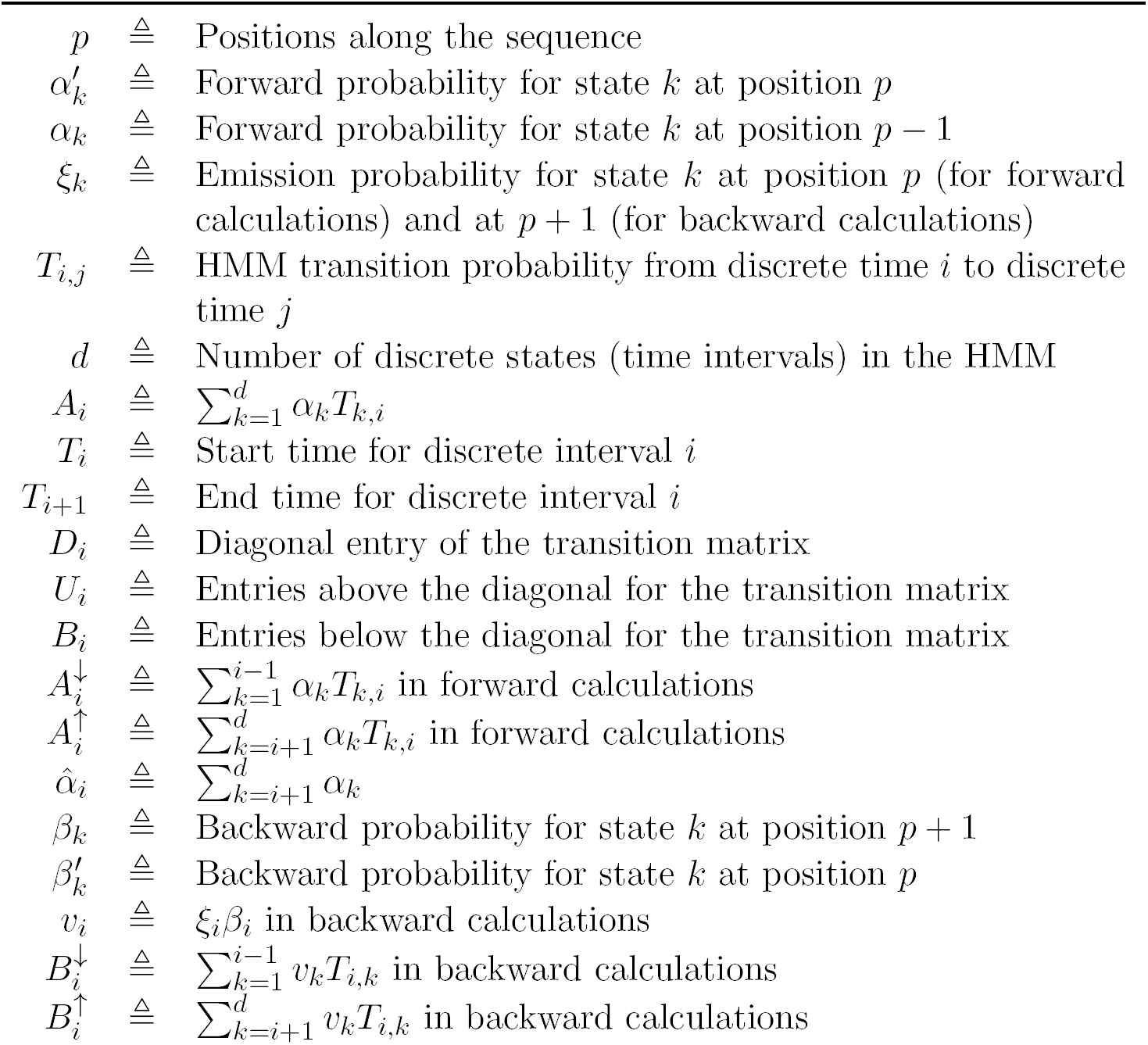
Table of notation for current section

##### 2.3.2.1 Forward probabilities

We want to compute 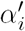, the forward probability at position *p* for state *i*, given a vector of forward probabilities for position *p* — 1 (which we denote as *α*_*k*_, dropping the position index to simplify notation). Using standard considerations from hidden Markov models, this can be obtained as 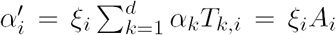, where ξ_*i*_ represents the emission probability for the observation at position *p* (dropped to simplify the notation) given state *i*. Because this operation involves a vector-matrix multiplication, the cost of computing 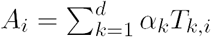 is linear in d, and because d forward probabilities need to be computed, the overall cost will be quadratic in d. However, we note that the entries below the diagonal in T are all identical, since *Q* (*t*|*s*) in Eq. 12 does not depend on s for *t < s.* Furthermore, the ratio of subsequent columns in the transition matrix can be computed as

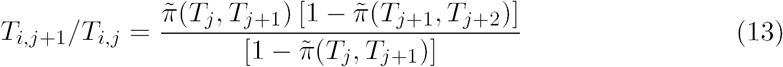

(see Appendix). This ratio does not depend on *i*, so that it will be the same for all rows of the *T* matrix, as long as the entries are above the diagonal. Taken together, these observations imply that the sum 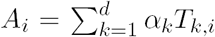 can be computed recursively in constant time. We assume the following quantities have been precomputed (in time linear in *d*), and are available for the computation of *A*_*i*_:

- The diagonal entries of the transition matrix *D*_*i*_ = *T*_*i,i*_ for *i* ∈ [1, *d*].
- The elements right above the diagonal *U*_*i*_ = *T*_*i*-1)*i*_, for *i* ∈ [2, *d*].
- The elements right below the diagonal *B*_*i*_ = *T*_*i*+ 1)*i*_ for *i* ∈ [1,*d* — 1].
- The cumulative sum of the o vector of forward probabilities from the previous position,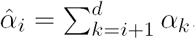.

We now rewrite the previous sum as

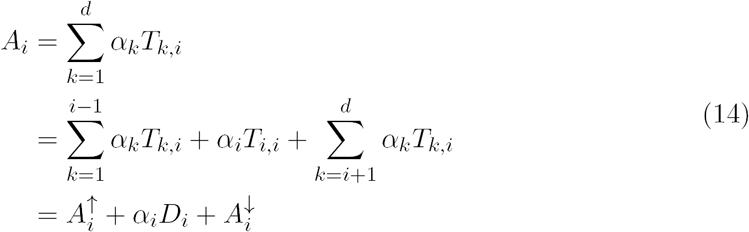

Then, the quantities 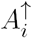 and 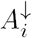 can be computed in constant time as follows:

- 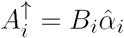
- 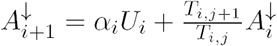, for *i* ∈ [2, *d*], after having set 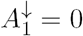

Having computed the above quantities (in time linear in d), all entries 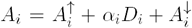 can be computed in linear time. The final forward vector is obtained multiplying the emission probabilities to obtain 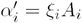.

##### 2.3.2.2 Backward probabilities

The linear-time backward calculations can be obtained in a similar way. In this case, given ξ, the emission probability vector at sequence position *p* + 1, and *β*, the backward probability vector for position *p* + 1, we want to compute 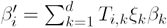 the backward probablity at state *i*, position *p*. We again use observations (1) and (2) from the previous section to efficiently compute this sum. It is convenient to define the vector *v* such that *v*_*i*_ = *ξ*_*i*_*β*_*i*_. As in the previous case, we rewrite the above sum as

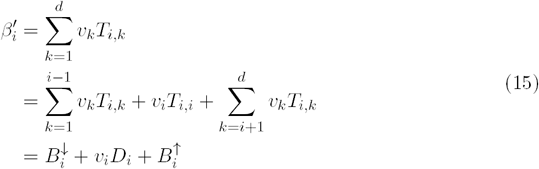

We have previously noted that the ratio of subsequent columns above the diagonal is constant (see Appendix). We now note that the same holds for the ratio of columns. In particular, it can be shown (see Appendix), that

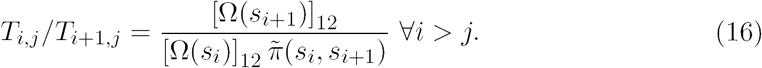

Using this result, the quantities 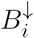 and 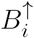 can be efficiently computed as

- 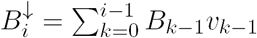, having set 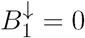.
- 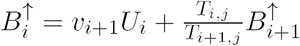, having set 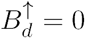.

From these quantities, we can then obtain 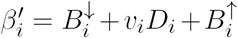. Note that these calculations hold for the SMC, SMC’ and CSC models, provided the corresponding transition matrices are used to compute entries of the Ω vector. Inhomegeneous (e.g. exponential) models can be handled by computing the corresponding coalescent quantities in the above calculations.

##### 2.3.2.3 Approximate decoding for stretches of identical observations

When ascertained data is analyzed and no information on the sequence content between observed *ℓ*_*p*_ markers is available, the linear time algorithm described above yields exact posterior TMRCA probabilities. Using a locus-skipping approximation, it is also possible to use the same linear-time forward-backward algorithm for the analysis of sequencing data, where we wish to obtain TMRCA estimates for *ℓ*_*p*_ loci (e.g. polymorphic loci), while accounting for the fact that all sites between any other two contiguous observations share the same emission probabilities (e.g. they are all monomorphic in the analyzed sample, or homozygous if frequency information is not used in the emission model). To this end we note that the forward step of the forward-backward algorithm between two sites separated by a stretch of *n* identical observations requires computing the product α′ = *α*(*TE*_*s*_)^*n*^*TE*_*p*_, where *T* is the transition matrix between two sites in the region, *E*_*s*_ is a diagonal matrix with the emission probability for a given emission character (e.g. homozygous/monomorphic site), and *E*_*p*_ is a diagonal matrix with emission for the site at position *p* in the sequence. We observe that, for relatively small genetic distances between the two observed sites, and for realistic demographic models, the matrix *T* is close to diagonal. Thus, we can use the commutative property of diagonal matrices to approximate the product (*TE*_*s*_)^*n*^ as 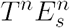. Having done that, we can now rely on the previosly described linear time algorithm to compute the product 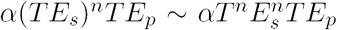. In the ASMC program, the matrices *T*^*n*^ and 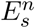 are precomputed (in linear time) and stored so that these need not be computed for each analyzed haploid pair. Note that the ASMC uses genetic distances from a human recombination map, rather than assuming a constant recombination rate along the genome, so that the matrix *T*^*n*^ will actually depend on genomic position, while the emission matrix 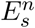 will only depend on the number of loci between a pair of sites.

## 3 Appendix

### 3.1 Ratio of columns in the transition matrix

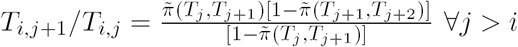 *Proof:*

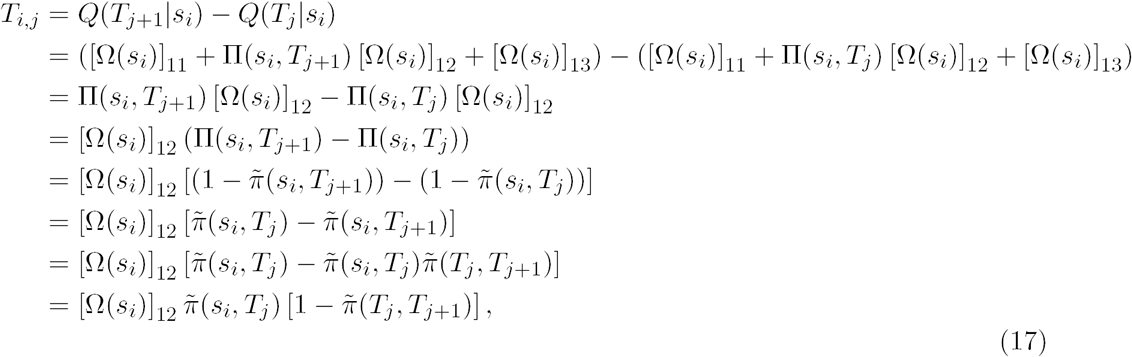

which implies

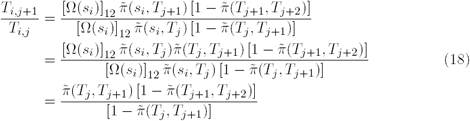

### 3.2 Ratio of rows in the transition matrix

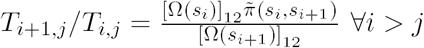 Again, using

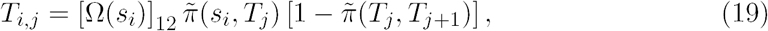

we have

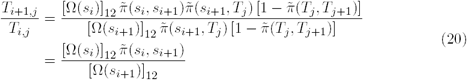

### 3.3 Above diagonal elements

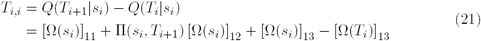

and

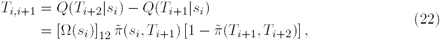

